# Mutations in enterobacterial common antigen biosynthesis restore outer membrane barrier function in *Escherichia coli tol-pal* mutants

**DOI:** 10.1101/480533

**Authors:** Xiang’Er Jiang, Wee Boon Tan, Rahul Shrivastava, Deborah Chwee San Seow, Swaine Lin Chen, Xue Li Guan, Shu-Sin Chng

## Abstract

The outer membrane (OM) is an essential component of the Gram-negative bacterial envelope that protects cells against external threats. To maintain a functional OM, cells require distinct mechanisms to ensure balance of proteins and lipids in the membrane. Mutations in OM biogenesis and/or homeostasis pathways often result in permeability defects, but how molecular changes in the OM affect barrier function is unclear. Here, we seek potential mechanism(s) that can alleviate permeability defects in *Escherichia coli* cells lacking the Tol-Pal complex, which accumulate excess PLs in the OM. We identify mutations in enterobacterial common antigen (ECA) biosynthesis that re-establish OM barrier function against large hydrophilic molecules, yet did not restore lipid homeostasis. Furthermore, we demonstrate that build-up of biosynthetic intermediates, but not loss of ECA itself, contributes to the rescue. This suppression of OM phenotypes is unrelated to known effects that accumulation of ECA intermediates have on the cell wall. Finally, we reveal that an unusual diacylglycerol pyrophosphoryl-linked lipid species also accumulates in ECA mutants, and might play a role in the rescue phenotype. Our work provides insights into how OM barrier function can be restored independent of lipid homeostasis, and highlights previously unappreciated effects of ECA-related species in OM biology.

## Introduction

Gram-negative bacteria are surrounded by a multilayered cell envelope consisting of the inner membrane (IM), the peptidoglycan layer, and the outer membrane (OM). This envelope structure, in particular the OM, plays an essential role in preventing toxic molecules from entering the cell, contributing to intrinsic resistance of Gram-negative bacteria against many antibiotics and detergents (Nikaido, 2003). The OM bilayer is asymmetric and has a unique lipid composition, comprising lipopolysaccharides (LPS) in the outer leaflet and phospholipids (PLs) in the inner leaflet. In the presence of divalent cations, LPS molecules in the outer leaflet pack together to form an impervious monolayer (Raetz & Whitfield, 2002); OM structure and lipid asymmetry are thus key determinants for its barrier function. Furthermore, the OM is essential for growth, highlighting the importance of understanding how it is established and maintained.

The biogenesis of the OM has been relatively well studied. Key components, including LPS, integral β-barrel proteins, and lipoproteins, are transported and assembled unidirectionally into the OM by the Lpt (Okuda et al., 2016), Bam (Hagan et al., 2011), and Lol machinery (Okuda & Tokuda, 2011), respectively. Recently, several systems, including the OmpC-Mla system and the Tol-Pal complex, have been implicated in PL transport between the IM and the OM (Shrivastava et al., 2017; Ekiert et al., 2017; Ercan et al., 2018; Thong et al., 2016; Hughes et al., 2019). However, much of bulk PL transport between the two membranes, which occurs in both directions (Donohue-Rolfe & Schaechter, 1980; Jones & Osborn, 1977; Langley et al., 1982), is still largely unknown (Shrivastava & Chng, 2019). Nonetheless, an intricate balance between these different OM biogenesis pathways needs to be maintained at all times to ensure a stable OM. Such homeostasis of OM components is critical for proper barrier function; yet, we do not fully understand the exact molecular mechanisms of how changes in the OM can translate to permeability defects. In many cases, mutations in OM biogenesis (e.g. *lptD4213, degP, bamB, tol-pal*) result in compromised OM barrier function against both hydrophobic molecules, such as detergents, and even large hydrophilic antibiotics, including vancomycin. One common problem with the OM in these mutants appears to be perturbations in the lipid asymmetry. This give rises to PL bilayer patches that would allow easier diffusive passage of hydrophobic compounds, but not hydrophilic ones. How large hydrophilic drugs like vancomycin cross the OM in mutants with defective OM biogenesis remains elusive.

The Tol-Pal complex is a conserved multi-protein system that forms a trans-envelope bridge across the cell envelope in Gram-negative bacteria, and has been known to be generally important for OM stability and integrity (Lloubes et al., 2001; Sturgis, 2001; Cascales et al., 2000). It comprises two sub-complexes: TolQRA in the IM and TolB-Pal at the OM. We recently demonstrated that the Tol-Pal complex is involved in the maintenance of OM lipid homeostasis in *Escherichia coli* (Shrivastava et al., 2017). Cells lacking the intact complex accumulate excess PLs in the OM, likely due to defects in retrograde (OM-to-IM) PL transport. This increase in PL content may contribute to many known OM permeability/stability phenotypes in *tol-pal* mutants, including hypersensitivity to antibiotics and detergents, leakage of periplasmic content, and OM hypervesiculation (Lloubes et al., 2001; Bernadac et al., 1998). The Tol-Pal complex also plays a key role in cell division; mutant strains lacking functional Tol-Pal exhibits division defects under extreme osmolarity due to incomplete septal cell wall separation and/or improper OM invagination (Gerding et al., 2007; Yakhnina & Bernhardt, 2020). Its role in cell division appears to be independent of its function in OM lipid homeostasis (Yakhnina & Bernhardt, 2020).

Lipid dyshomeostasis in cells lacking the Tol-Pal complex affects OM barrier function in a way that allows entry of both hydrophobic molecules and large hydrophilic antibiotics. To gain insight(s) into OM lipid homeostasis and how it possibly affects OM function, we asked whether cells lacking the Tol-Pal complex could restore OM barrier function by compensatory mutations in other pathways. We found that disruption in the biosynthesis of enterobacterial common antigen (ECA), a cell surface polysaccharide in the Enterobacteriaceae family (Kuhn et al., 1988), rescues OM permeability defects in *tol-pal* mutants, especially against large hydrophilic molecules including antibiotics and periplasmic proteins. Interestingly, in these suppressor mutants, OM lipid homeostasis is still defective even though barrier function is restored. We demonstrated that the rescue requires the accumulation of ECA intermediates, yet is independent of known effects of such build-up on cell wall integrity. Finally, we detected unusual ECA-related lipids in these cells, raising the possibility that they might play a role in restoring OM phenotypes. Our work reveals that the OM barrier, specifically against large hydrophilics, can somehow be re-established even during lipid dyshomeostasis, highlighting the complex and dynamic nature of bacterial OM permeability.

## Results

### Loss-of-function mutations in ECA biosynthesis rescue OM permeability defects in *tol-pal* mutants

Cells lacking the Tol-Pal complex have compromised OM integrity due to the accumulation of excess PLs (Shrivastava et al., 2017). To gain insight(s) into this phenomenon, we sought to isolate genetic suppressor mutations that could restore OM barrier function in a Δ*tol-pal* mutant. We selected for suppressors that rescue sensitivity to vancomycin, the large cell wall-targeting antibiotic that cannot penetrate an intact and functional OM (Krishnamoorthy et al., 2016). Cells have no intrinsic mechanism to alter cell wall structure to give rise to resistance against vancomycin; consistently, suppressor colonies appeared at a low frequency of ∼1 × 10^−9^. We obtained 36 individual suppressor strains, all of which exhibited stable resistance phenotypes against vancomycin similar to a wild-type (WT) strain. We further examined the susceptibilities of these strains against other commonly used antibiotics, including rifampicin and erythromycin. An intact OM also impedes the entry of rifampicin and erythromycin but less effectively than against vancomycin (Krishnamoorthy et al., 2016). Eight of the suppressors exhibited near wild-type level resistance against vancomycin and erythromycin (Fig. S1A), thus appearing to have restored OM barrier function. Consistent with this, many of these strains release less periplasmic RNase I into the media during growth when compared to the parent Δ*tol-pal* mutant (Fig. S1B). We sequenced the genomes of the parent strain and all eight suppressors. Each suppressor strain contains multiple mutations relative to the parent Δ*tol-pal* mutant (Table S1). Many strains contain mutations in either *pgm* or *mdoH* (*opgH*), which have already been implicated in conferring vancomycin resistance, especially at low temperature (Stokes et al., 2016). Interestingly, all of these strains have mutations in genes involved in the biosynthesis of ECA (*wecB, wecC*, or *wecF*), so we decided to further investigate this genetic interaction.

Six strains contain mutations in *wecC*, which encodes a dehydrogenase enzyme in the ECA synthesis pathway (Kuhn et al., 1988; Meier-Dieter et al., 1990). A 6-bp in-frame insertion in *wecC*, termed *wecC\**, is common to four of these strains, suggesting that this allele may be important for restoring OM barrier function in the Δ*tol-pal* mutant. To validate this, we re-constructed the *wecC\** mutation in the native locus using a negative selection technique (Khetrapal et al., 2015) and confirmed that this allele alone is able to partially rescue antibiotic sensitivity (Figs. 1A, S2A) and periplasmic leakiness, albeit to different extents in Δ*tolQ*, Δ*tolA*, and Δ*tolB* strains (Fig. S2B). These partial rescue phenotypes are consistent with the idea that *pgm* and/or *mdoH* (*opgH*) mutations found in the original suppressor strains also contribute to restoring OM function (Stokes et al., 2016).

**Fig. 1.**
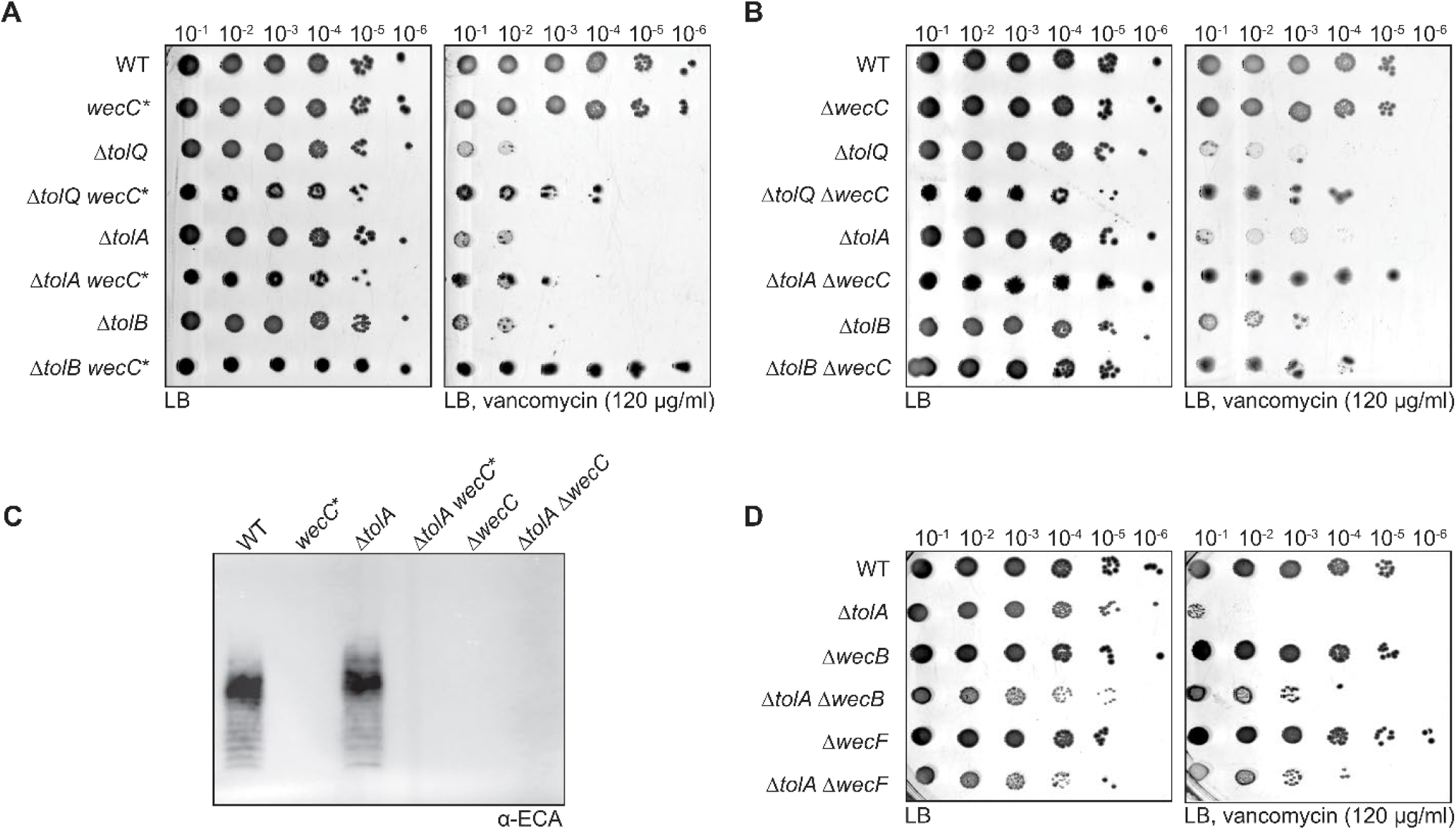
Loss-of-function mutations in the ECA pathway rescue vancomycin sensitivity in *tol-pal* mutants. Vancomycin sensitivity of indicated *tol-pal* strains with or without (A) *wecC*\*, (B) Δ*wecC*, and (D) Δ*wecB*/Δ*wecF* mutations based on efficiency of plating (EOP) on LB agar plates supplemented with vancomycin (120 μg/ml). (C) ECA levels in wild-type (WT) and Δ*tolA* strains with or without *wecC*\* and Δ*wecC* mutations as judged by immunoblot analysis using α-ECA antibody. Samples were normalized by OD_600_ and treated with proteinase K prior to Tricine SDS-PAGE/immunoblotting.

The *wecC\** mutation results in the insertion of two amino acids (Pro and Gly) five residues away from the predicted active site Cys in the full-length protein, suggesting that WecC function may be disrupted. Consistent with this, we did not detect any ECA in strains containing this *wecC\** allele (Fig. 1C). Furthermore, we showed that deletion of *wecC* also partially restores vancomycin resistance in *tol-pal* strains (Figs. 1B and C). We isolated *wecB* and *wecF* mutations in two of the sequenced suppressor strains (Table S1); we therefore constructed Δ*wecB* and Δ*wecF* mutants to test their rescue phenotypes. Both null mutations partially rescue vancomycin sensitivity in the Δ*tolA* strain (Fig. 1D). Finally, we showed that the Δ*wecC* mutation does not suppress vancomycin sensitivity in strains with other OM defects (*lptD4213* and *bepA*) (Fig. S3). We conclude that loss-of-function mutations in ECA biosynthesis likely restore the OM barrier function specifically in strains lacking the Tol-Pal complex.

### Mutations in ECA biosynthesis restore OM barrier function in *tol-pal* strains independent of Rcs phosphorelay pathway and/or capsular polysaccharide biosynthesis

Mutations in the ECA pathway are known to trigger the Rcs phosphorelay stress response (Castelli & Vescovi, 2011). *tol-pal* mutations also strongly activate the Rcs signaling cascade (Clavel et al., 1996). Consequently, combined *tol-pal/*ECA mutant colonies exhibit strongly mucoidal phenotypes, presumably due to the over-production of capsular polysaccharides (colanic acids), which is regulated by the Rcs pathway (Stout & Gottesman, 1990; Trisler & Gottesman, 1984). We therefore considered whether hyperactivation of the Rcs stress response or up-regulation of colanic acid biosynthesis contributed to the suppression of vancomycin sensitivity by ECA mutations in the *tol-pal* strains. To test this idea, we examined vancomycin sensitivity in *rcsC* (encoding the histidine sensor kinase of the Rcs pathway) (Stout & Gottesman, 1990) or *wcaJ* (encoding the glycosyl transferase that initiates colanic acid biosynthesis) (Stevenson et al., 1996) mutants. Deleting *rcsC* or *wcaJ* did not prevent the Δ*wecC* mutation from suppressing vancomycin sensitivity in the Δ*tolA* strain; instead, the Δ*rcsC* Δ*tolA* Δ*wecC* and Δ*wcaJ* Δ*tolA* Δ*wecC* mutants display full resistance against vancomycin, similar to WT cells (Fig. 2A). The mechanism by which the Δ*wecC* mutation suppresses vancomycin sensitivity in *tol-pal* strains is therefore independent of the Rcs phosphorelay and/or colanic acid synthesis.

**Fig. 2.**
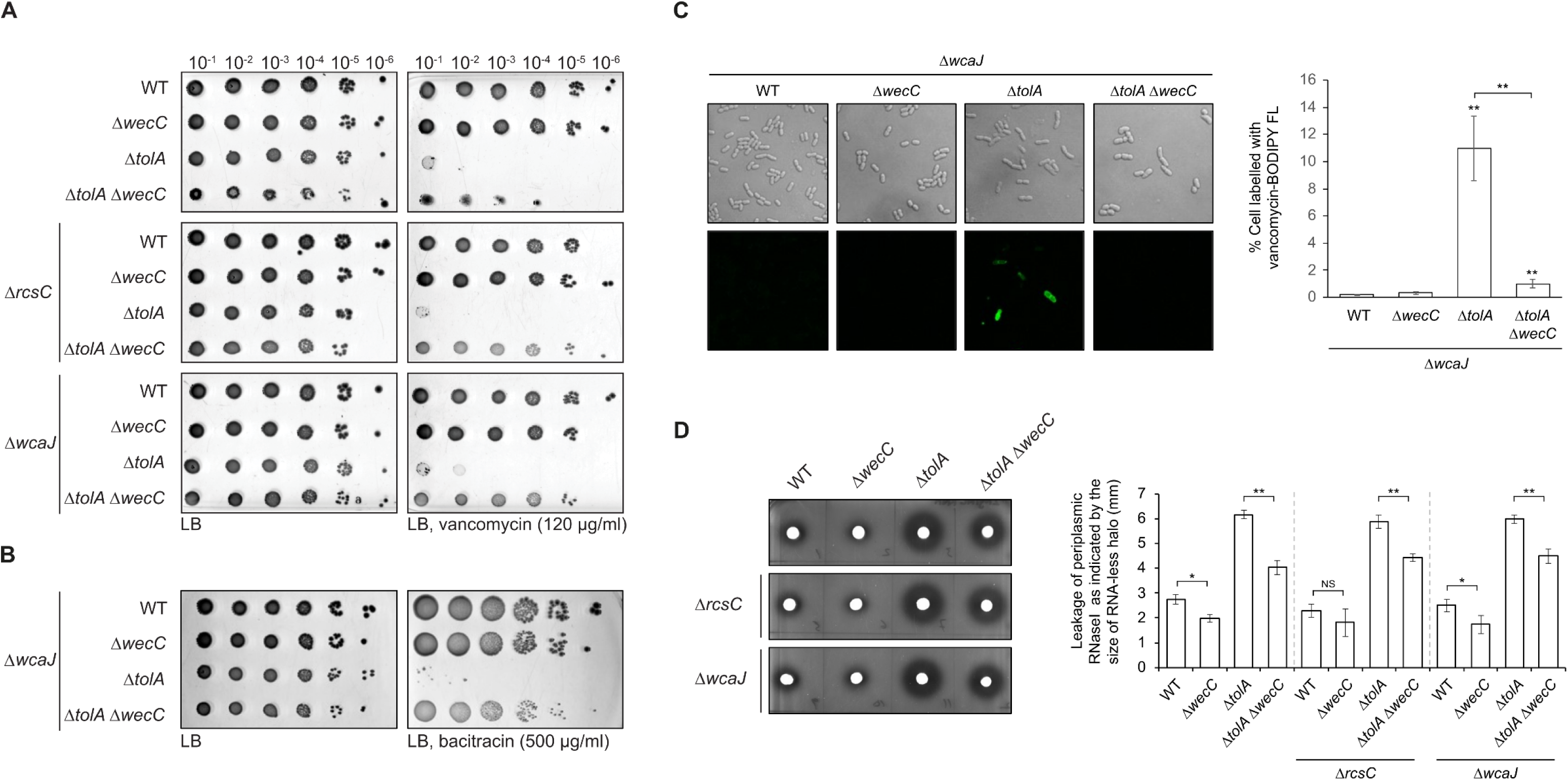
Restoration of OM permeability defects by Δ*wecC* in cells lacking TolA does not require the Rcs phosphorelay cascade and/or capsular polysaccharide biosynthesis. (A) Vancomycin sensitivity of indicated strains in either Δ*rcsC* or Δ*wcaJ* backgrounds based on EOP on LB agar plates supplemented with vancomycin (120 μg/ml). (B) Bacitracin sensitivity of indicated strains in Δ*wcaJ* backgrounds based on EOP on LB agar plates supplemented with bacitracin (500 μg/ml, ≥60 U/mg). (C) Confocal microscopy images of indicated strains (Δ*wcaJ* background) after incubation with BODIPY FL vancomycin. Quantification of the percentage of cells circumferentially labelled with BODIPY FL vancomycin from different fields of view (n = 6) is shown on the right; error bars represent standard deviation. Student’s t-tests: **, *p* < 0.005 (as compared to WT, unless otherwise indicated). (D) RNase I leakage in the same strains as (A), as judged by RNA degradation (halo formation) around cells spotted on LB agar plates containing yeast RNA, subsequently precipitated with trichloroacetic acid. Quantification of the distances between the edges of the macrocolony and the halo (n = 3) is shown on the right; error bars represent standard deviation. Student’s t-tests: *, p < 0.05; **, p < 0.005.

To further demonstrate that the OM barrier function was indeed restored, we showed that Δ*wecC* partially rescues rifampicin and erythromycin sensitivity (Fig. S4) and fully restores bacitracin resistance in the Δ*wcaJ* Δ*tolA* strain (Fig. 2B). Furthermore, we demonstrated that a fluorescent analog of vancomycin (BODIPY FL vancomycin) could only penetrate the OM and label peptidoglycan in the Δ*wcaJ* Δ*tolA* strain, but not the Δ*wcaJ* Δ*tolA* Δ*wecC* triple mutant (Fig. 2C). The Δ*wcaJ* Δ*tolA* Δ*wecC* strain also exhibited reduced periplasmic leakiness compared to the Δ*wcaJ* Δ*tolA* mutant (Fig. 2D). Rifampicin (MW ∼823) and erythromycin (MW ∼734) are much smaller than vancomycin (MW ∼1449) and bacitracin (MW ∼1423). We conclude that the Δ*wecC* mutation restores OM barrier function in the *tol-pal* strains lacking colanic acid synthesis, especially against the passage of larger hydrophilic molecules, including vancomycin, bacitracin, and periplasmic proteins (e.g. RNase I).

We have observed that the Δ*wecC* mutation rescues vancomycin sensitivity in the Δ*tolA* strain; however, we noted that different transductants display varying extents of suppression (Fig. S5). We found that removing RcsC or WcaJ greatly improved the consistency of suppression of the Δ*tolA* phenotype by the Δ*wecC* mutation. This suggests that the initial variability in suppression could be due to varying mucoidal phenotypes, possibly because of stochasticity of colanic acid overproduction. This might additionally explain why the *wecC*\* and Δ*wecC* mutations also showed different levels of rescue in Δ*tolQ*, Δ*tolA*, and Δ*tolB* strains that still produce colanic acids (Figs. 1, S2). To eliminate this inconsistency, we used strains deleted of *wcaJ* for the rest of this study.

### Mutations in ECA biosynthesis rescue OM barrier function without restoring lipid homeostasis in *tol-pal* mutants

Cells lacking the Tol-Pal complex accumulate excess PLs (relative to LPS) in the OM due to defective retrograde PL transport (Shrivastava et al., 2017). Since mutations in the ECA biosynthetic pathway rescue OM defects in *tol-pal* mutants (Fig. 1), we hypothesized that OM lipid homeostasis is restored in these strains. To test this idea, we examined steady state OM lipid compositions in Δ*wcaJ* Δ*tolA* Δ*wecC* mutant cells by measuring the distribution of [^32^P]-phosphate-labelled PLs between the IM and the OM, and also determining the ratio of PLs to LPS (also labelled with [^32^P]) in the OM. Consistent with our previous findings, the Δ*tolA* mutant contained more PLs in the OM than WT cells (here in the Δ*wcaJ* background). Specifically, cells lacking TolA accumulate ∼1.7-fold excess PLs in the OM (relative to the IM) (Fig. 3A). Furthermore, these cells have ∼1.4-fold higher PL/LPS ratio in the OM (Fig. 3B). To our surprise, deleting *wecC* in the Δ*tolA* strain neither re-established intermembrane PL distribution (Fig. 3A), nor restored the OM PL/LPS ratio back to wild-type (Fig. 3B). Consistently, the Δ*tolA* and Δ*tolA* Δ*wecC* strains produced similarly high levels of OM vesicles (Fig. 3C), a phenotype in part attributed to excess PLs in the OM (Shrivastava et al., 2017). Even though the OM barrier function (restriction of vancomycin/bacitracin entry and periplasmic RNase I release) has largely been restored (Fig. 2), we conclude that mutations in ECA biosynthesis do not restore OM lipid homeostasis in cells lacking the Tol-Pal complex.

**Fig. 3.**
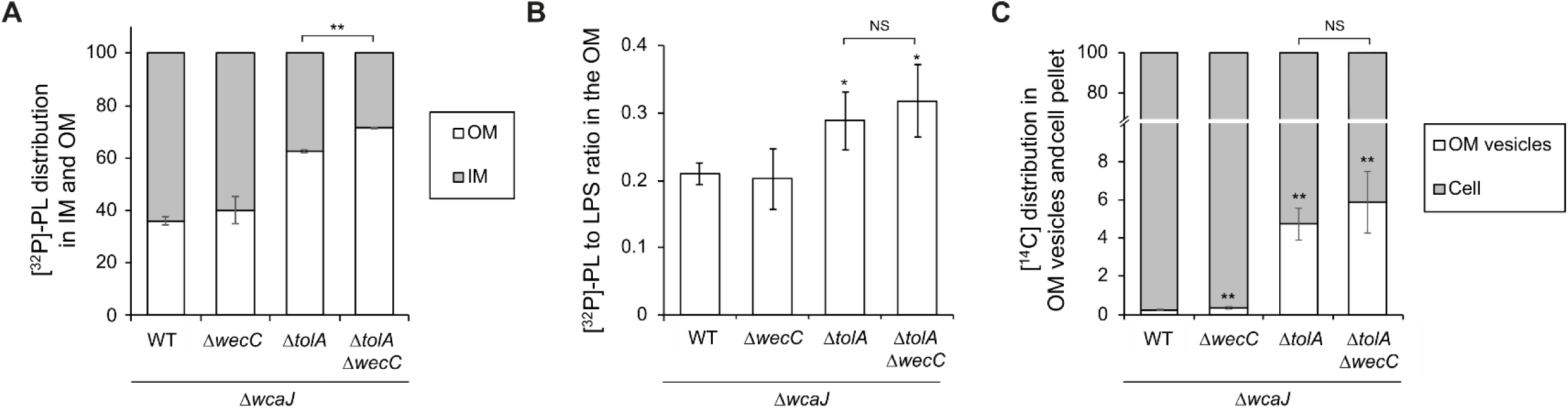
Δ*wecC* mutation does not restore OM lipid homeostasis in cells lacking TolA. (A) Steady-state [^32^P]-labelled PL distribution profiles of indicated Δ*wcaJ* strains. Cells were grown in the presence of [^32^P]-phosphate to label PLs in the IMs and OMs. [^32^P]-scintillation counts detected in PLs extracted from IM and OM fractions were expressed as a percentage of their sums, averaged across three replicate experiments. (B) Steady-state PL:LPS ([^32^P]-phosphate labelled) ratios in the OMs of indicated Δ*wcaJ* strains. Arbitrary OM PL:LPS ratios were calculated based on [^32^P]-scintillation counts detected from PL or LPS, differentially extracted from the OM fractions of the indicated strains. (C) Steady-state distribution of [^14^C]-acetate-labelled lipids found associated with cells (total membranes) or OM vesicles for the indicated strains. Error bars represent standard deviations calculated from triplicate experiments. Student’s t-tests: *, *p* < 0.05; **, *p* < 0.005; NS, not significant (as compared to WT, unless otherwise indicated).

### Accumulation of ECA intermediates along the biosynthetic pathway is critical for rescue of OM barrier function in *tol-pal* strains

The initial steps of ECA biosynthesis involve successive addition of three sugar moieties (*N*-acetyl-D-glucosamine (GlcNAc), *N*-acetyl-D-mannosaminuronic acid (ManNAcA), and 4-acetamido-4,6-dideoxy-D-galactose (Fuc4NAc)) to the undecaprenyl phosphate (und-P) lipid carrier to form Lipid III^ECA^ (Kuhn et al., 1988; Rahman et al., 2001). After its synthesis at the inner leaflet of the IM, Lipid III^ECA^ is flipped across the membrane by WzxE (Rick et al., 2003). This trisaccharide repeating unit is then polymerized by WzyE to form the complete ECA polymer, whose polysaccharide chain length is regulated by WzzE (Kajimura et al., 2005; Barr et al., 1999). The whole ECA polymer, which is now carried on undecaprenyl pyrophosphate (und-PP), is finally transferred to form a phosphatidyl-linked species (ECA_PG_) before being transported to the OM (Kuhn et al., 1983; Rinno et al., 1980; Acker et al., 1986) (Fig. 4A). The enzymes mediating the last two steps are not known. Of note, two other minor forms of ECA are produced, including a soluble cyclic form in the periplasm (ECA_CYC_) and one that is found attached to LPS (ECA_LPS_) (Kuhn et al., 1988). The production of these two ECA forms is dependent on WzzE and the LPS O-antigen ligase WaaL, respectively (Kajimura et al., 2005; Schmidt et al., 1976; Mitchell et al., 2018).

**Fig. 4.**
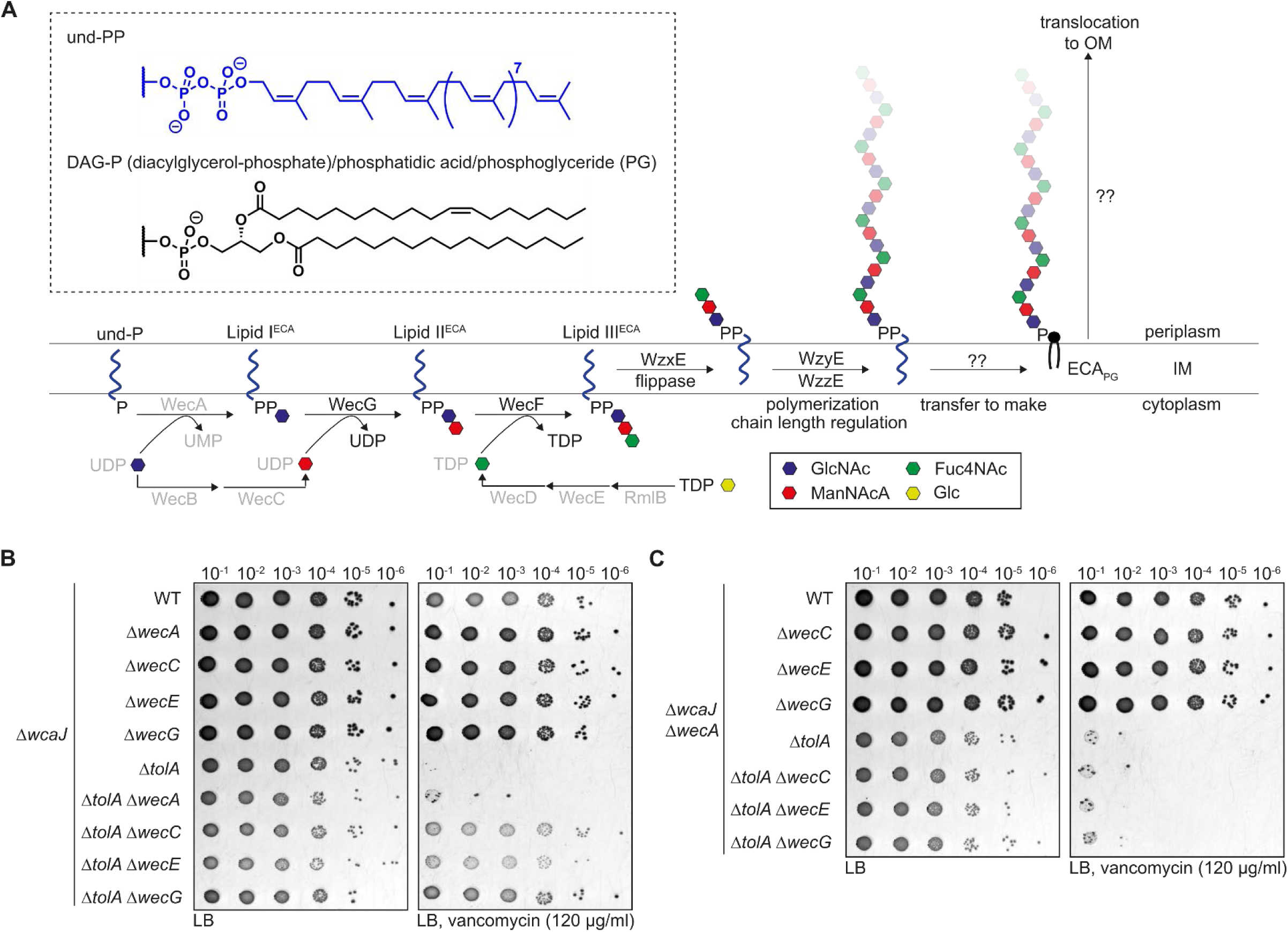
Accumulation of ECA biosynthetic intermediates restores vancomycin resistance in cells lacking TolA. (A) Schematic of the ECA biosynthetic pathway illustrating und-PP-GlcNAc-ManNAcA-Fuc4NAc (Lipid III^ECA^) synthesis at the inner leaflet of IM, and subsequent polymerization at the outer leaflet. The final ECA polymer (ECA_PG_) is attached to a phosphoglyceride (i.e. diacylglycerol-phosphate (DAG-P) aka phosphatidic acid) and transported to the OM. Chemical structures of und-PP and DAG-P anchors are shown. (B, C) Vancomycin sensitivity of the Δ*wcaJ* Δ*tolA* strain with or without indicated *wec* mutations in otherwise (B) WT or (C) Δ*wecA* backgrounds, based on efficiency of plating (EOP) on LB agar plates supplemented with vancomycin (120 ug/ml).

We have shown that loss-of-function mutations in *wecB, wecC*, and *wecF* rescue vancomycin sensitivity in *tol-pal* mutants (Fig. 1). Mutations in ECA biosynthesis result in both the loss of surface ECA and the build-up of intermediates (Lipid I/II/III^ECA^) along the pathway (Fig. 4A). To test whether ECA loss is important, we mutated *wecA*, which encodes the enzyme that catalyzes the first committed step in ECA biosynthesis. Interestingly, we found that the Δ*tolA* Δ*wecA* mutant (in the Δ*wcaJ* background) is equally sensitive to vancomycin as the Δ*tolA* mutant (Fig. 4B), suggesting that accumulation of intermediates along the ECA pathway, but not the loss of any form of ECA (ECA_PG_, ECA_LPS_, or ECA_CYC_), is responsible for the suppression. Consistent with this idea, removing other ECA biosynthetic enzymes that result in accumulation of intermediates (Lipid I^ECA^ in Δ*wecG*, or Lipid II^ECA^ in Δ*wecE*) fully rescues vancomycin sensitivity in the Δ*tolA* mutant (Fig. 4B). Expressing the deleted *wec* genes *in trans* reversed this effect (Fig. S6A). Importantly, rescue of vancomycin sensitivity in these strains was completely abolished when *wecA* was subsequently deleted (Fig. 4C). Likewise, expressing *wecA* in these strains *in trans* re-establish the rescue (Fig. S6B). We conclude that the build-up of Lipid I^ECA^ or Lipid II^ECA^ intermediates can somehow restore OM barrier function in the absence of the Tol-Pal complex.

We next tried to test whether mutations in later steps of ECA biosynthesis also suppress *tol-pal* phenotypes. We found that removing WzxE, the Lipid III^ECA^ flippase, does not rescue vancomycin sensitivity in the Δ*tolA* strain (Fig. S7A). However, the Δ*wzxE* mutant still produces full length ECA (Fig. S7B), presumably because the O-antigen flippase WzxB can also transport Lipid III^ECA^ (Rick et al., 2003), thus preventing its accumulation. Nonetheless, it has been reported that completely blocking translocation across the IM or subsequent polymerization of Lipid III^ECA^ causes toxicity in cells (Rick et al., 2003; Kajimura et al., 2005; Baba et al., 2006); removing both flippases (WzxE and WzxB) or the polymerase (WzyE) is lethal (Figs. S7C, D), precluding further analysis. We therefore turned our attention to WzzE. Remarkably, deletion of *wzzE* partially rescues vancomycin sensitivity in the Δ*tolA* mutant, and this phenotype is dependent on the presence of WecA (Fig. 5A). Cells lacking WzzE are known to lose modality in the ECA polymer, giving rise to a more random distribution of chain lengths (Barr et al. 1999). We validated this observation; there are relatively higher levels of shorter chain ECA, presumably including Lipid III^ECA^ precursors, in the Δ*wzzE* mutant (Fig. 5B). Taken together, our results suggest that build-up of short-chain or Lipid III^ECA^ intermediates can also possibly rescue OM defects in cells lacking the Tol-Pal complex.

**Fig. 5.**
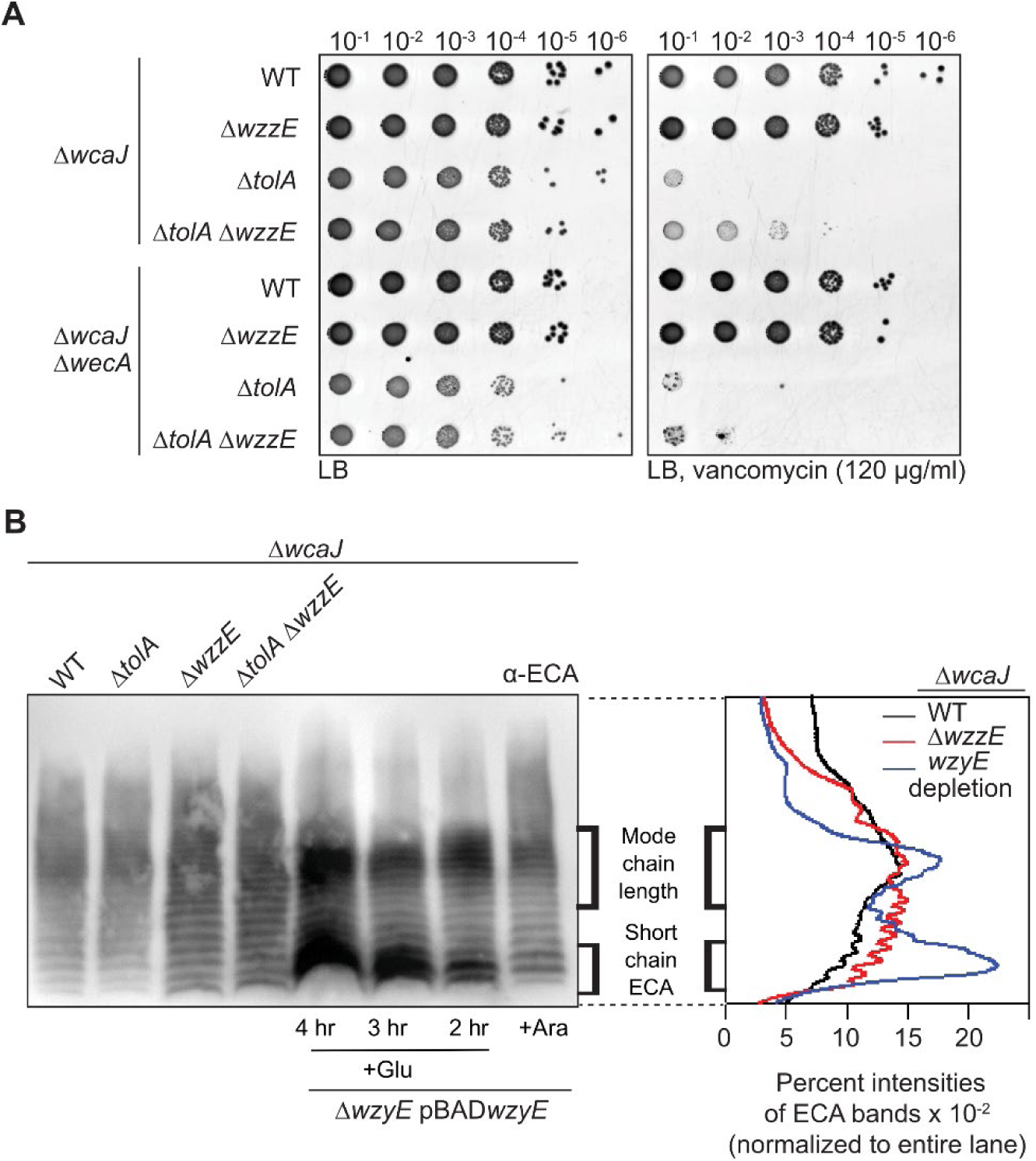
Partial accumulation of Lipid III^ECA^ and/or short chain ECA rescues vancomycin sensitivity in cells lacking TolA. (A) Vancomycin sensitivity of the Δ*wcaJ* Δ*tolA* strain with or without Δ*wzzE* mutation in otherwise WT or Δ*wecA* backgrounds, based on efficiency of plating (EOP) on LB agar plates supplemented with vancomycin (120 ug/ml). (B) ECA profiles in the indicated Δ*wcaJ* strains as judged by immunoblot analysis using α-ECA antibody. Samples were normalized by OD_600_ and treated with proteinase K prior to Tricine SDS-PAGE/immunoblotting. Profiles in strains progressively depleted of *wzyE* (at 2, 3, or 4 hours post subculture into media containing 0.2% glucose) are used to visualize accumulation of Lipid III^ECA^. Relative band intensities from ECA profiles of WT, Δ*wzzE*, and *wzyE-*depleted (for 3 hr) strains are shown on the right.

It has previously been reported that the terminal GlcNAc moiety attached to the *E. coli* K-12 LPS core oligosaccharide is derived from und-PP-GlcNAc (i.e. Lipid I^ECA^) via the action of WaaL (Feldman et al., 1999; Ruan et al., 2018). Since WaaL accepts different und-PP-linked substrates, it may be possible that corresponding modifications of LPS can occur in strains accumulating Lipid II/III^ECA^; these modified LPS might contribute to restored OM barrier function in the *tol*-*pal* mutants. To test this idea, we checked if the rescue of vancomycin sensitivity in the Δ*tolA* Δ*wecC* strain is dependent on WaaL. We found that deleting *waaL* did not confer vancomycin sensitivity to the Δ*tolA* Δ*wecC* mutant (Fig. S8), suggesting that the suppression of OM permeability defects is not due to WaaL-dependent LPS modifications.

### Suppression of OM permeability defects in *tol-pal* strains via accumulation of ECA intermediates is independent of und-P sequestration or σ^E^ and Cpx stress response pathways

Recently, it has been reported that the build-up of ECA “dead-end” intermediates can lead to sequestration of und-P, the common precursor for many sugar polymers in the cell envelope including peptidoglycan; this gives rise to shape defects such as filamentation and swelling (Jorgenson et al., 2016; Campos et al., 2018). In our strains that lack colanic acids, we also observe similar shape defects. We measured primarily larger cell width in the Δ*wecC* mutant, which were exacerbated in the Δ*tolA* background (Figs. S9A, B). The morphological changes observed in the Δ*tolA* Δ*wecC* strain are thus a combination of defects due to the loss of Tol-Pal function (Rassam et al., 2018), and und-P sequestration caused by accumulation of ECA intermediates (Jorgenson et al., 2016). To test whether und-P sequestration contributes to the rescue of vancomycin sensitivity in the Δ*tolA* Δ*wecC* strain, we examined if overexpression of MurA, which can alleviate und-P sequestration effects by directing the lipid carrier towards peptidoglycan biosynthesis (Jorgenson et al., 2016), could reverse the phenotypes. As expected, MurA overexpression was able to fully and partially revert the cell morphological defects in Δ*wecC* and Δ*tolA* Δ*wecC* strains, respectively (Figs. S9A, B, D). However, doing so did not alter vancomycin resistance in the Δ*tolA* Δ*wecC* mutant (Figs. S9C, D). Similarly, vancomycin sensitivity was not restored when overexpressing UppS, which is expected to increase the und-P pool (Jorgenson et al., 2016). These data indicate that the effect of accumulation of ECA intermediates in restoring OM barrier function in *tol-pal* mutants is independent of und-P sequestration.

Accumulation of ECA intermediates, specifically Lipid II^ECA^, is also known to stimulate the σ^E^ and Cpx stress response pathways (Danese et al., 1998). However, we did not observe significant induction or increased stimulation of σ^E^ when ECA intermediates were accumulated in WT or Δ*tolA* strains, respectively (Fig. S10A). Removing the Cpx pathway also did not affect vancomycin resistance in the Δ*tolA* Δ*wzzE* strain (Fig. S10B). We conclude that both σ^E^ and Cpx stress responses are also not involved in the ability of accumulated ECA intermediates to restore the OM barrier function in *tol-pal* mutants.

### Diacylglycerol pyrophosphoryl-linked species accumulate in ECA biosynthesis mutants in addition to undecaprenyl pyrophosphoryl-linked intermediates

We have shown that accumulation of ECA intermediates rescues vancomycin sensitivity in strains lacking the Tol-Pal complex. However, und-PP-linked intermediates (Lipid I/II/III^ECA^) may not be the only species accumulated in ECA biosynthesis mutants. In a *Salmonella* Typhimurium Δ*rmlA* mutant, which accumulates Lipid II^ECA^, a novel diacylglycerol pyrophosphoryl (DAG-PP)-linked species containing the first two sugars of ECA (GlcNAc and ManNAcA) was detected at comparable levels (Rick et al., 1998). We therefore sought to determine whether DAG-PP-linked adducts could also be found in our *E. coli* ECA mutants. Using high resolution mass spectrometry (MS), we analyzed lipids extracted from cells lacking WecG; as expected, we saw accumulation of the und-PP-GlcNAc or Lipid I^ECA^ (m/z 1128.7026) intermediate (Fig. 6). Furthermore, we demonstrated that DAG-PP-GlcNAc species are indeed present in these cells, but not in WT (Figs. 7A, B). Specifically, we detected peaks with m/z values corresponding to DAG-PP-GlcNAc species with various fatty acid compositions in the DAG moiety, namely 32:1 (m/z 928.4927), 34:1 (m/z 956.5243), 34:2 (m/z 954.5076), and 36:2 (m/z 982.5391); these are in fact the major DAGs found in native PLs in *E. coli* (Oursel et al., 2018). Chemical structures of the 32:1 and 34:1 species were assigned and elucidated based on fragmentation patterns in MS/MS (Fig. 8). Furthermore, we showed that the same species were specifically found in the IM but not the OM of the Δ*wecG* mutant, as well as the Δ*wecC* strain (Fig. 7C). It is worth noting that DAG-PP has one extra phosphate moiety and is therefore structurally distinct from phosphatidic acid (i.e. diacylglycerol monophosphate or DAG-P), the final lipid carrier of ECA; instead, the DAG-PP-GlcNAc and DAG-PP-GlcNAc-ManNAc species (Rick et al., 1998) have been suggested as structural precursors to the MPIase glycolipid (Nishiyama et al., 2012; Sawasato et al., 2019a), which is required for protein integration and translocation at the IM. How these DAG-PP-linked species are generated is not fully understood, but their existence highlights the need to consider possible effects of these unusual lipids in our ECA biosynthesis mutants, especially in the context of rescuing OM barrier function in *tol-pal* strains.

**Fig. 6.**
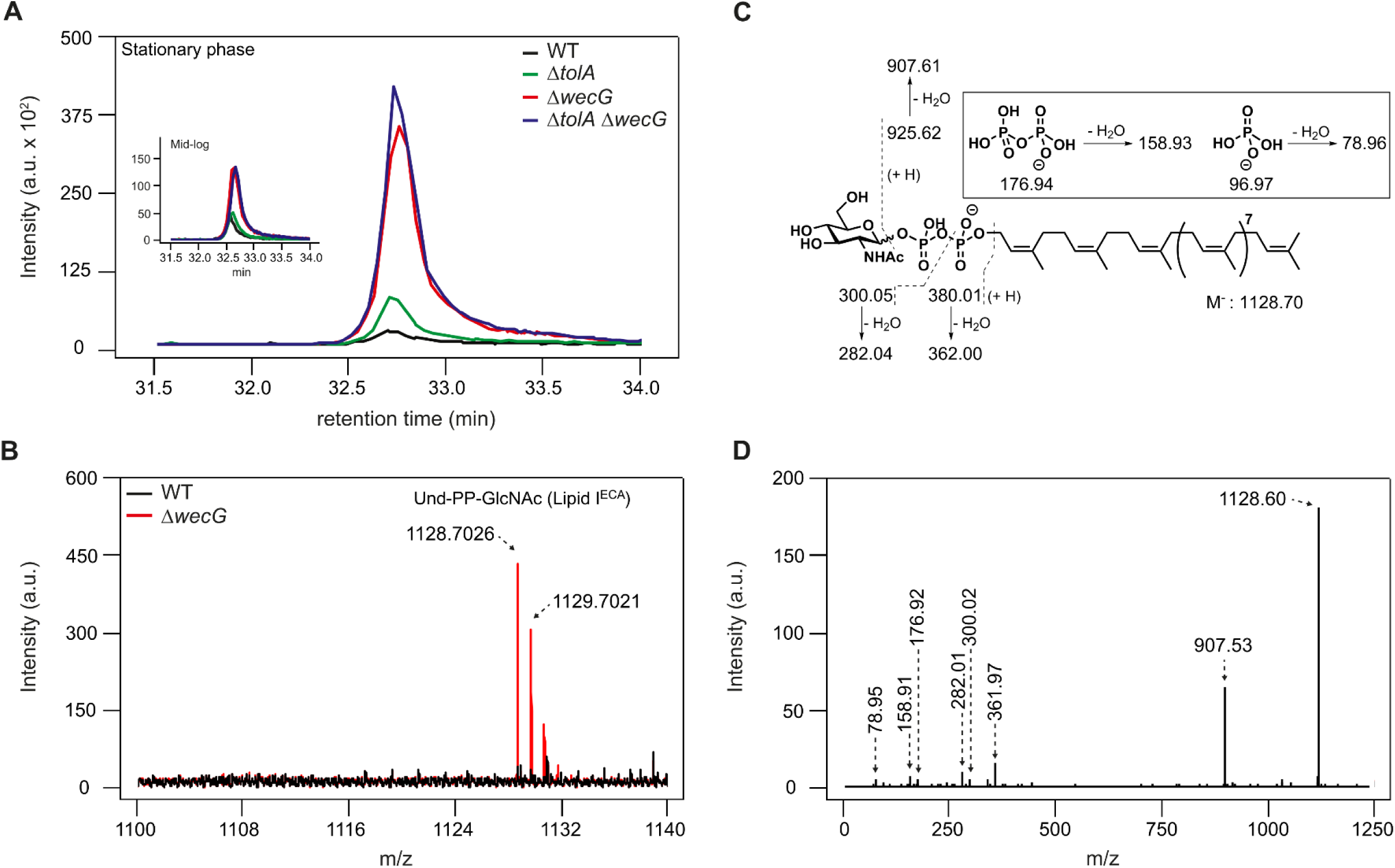
Strains lacking WecG accumulate und-PP-GlcNAc (Lipid I^ECA^) intermediates. (A) Ion chromatograms (XICs) of und-PP-GlcNAc (m/z 1128.7026) extracted from LC-MS analyses of total lipids isolated from indicated Δ*wcaJ* strains grown to stationary phase. Inset, corresponding XICs for the same lipid in strains grown to mid-logorithmic phase. (B) Mass spectra obtained from integration of XIC peak region in (A) of Δ*wcaJ* (WT) and Δ*wcaJ* Δ*wecG* strains. WT peaks (black) are overlaid on top of Δ*wecG* peaks (red), illustrating strong und-PP-GlcNAc signals in the latter strain. (C) Proposed fragmentation pattern with mass assignments, and (D) MS/MS spectrum at low collision energy (−55V) of the und-PP-GlcNAc species.

**Fig. 7.**
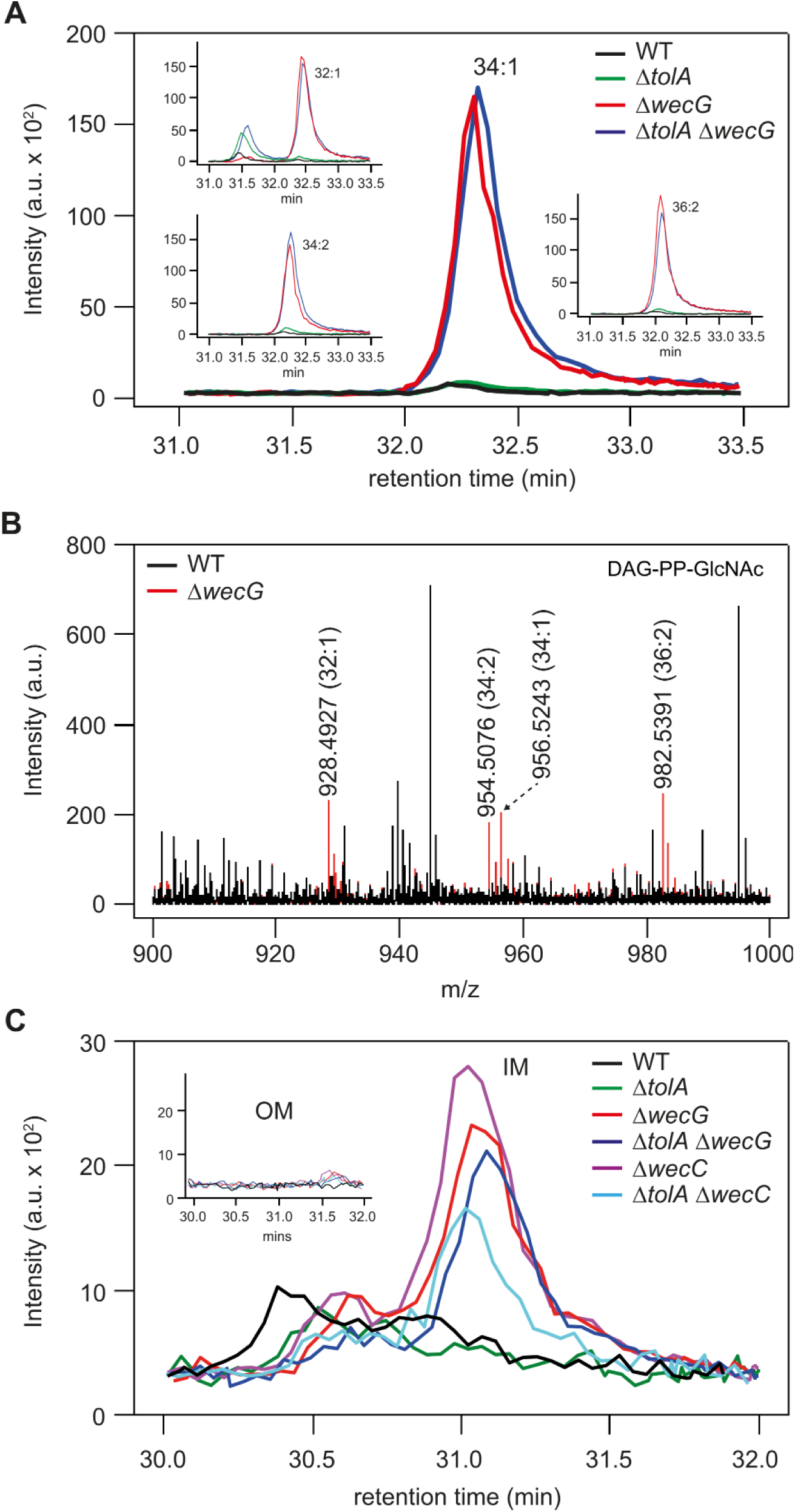
Strains lacking WecG or WecC accumulate DAG-PP-GlcNAc species in the IM. (A) Ion chromatograms (XICs) of 34:1 DAG-PP-GlcNAc (m/z 956.5243) extracted from LC-MS analyses of total lipids isolated from indicated Δ*wcaJ* strains grown to mid-logarithmic phase. Inset, XICs of corresponding 32:1 (m/z 928.4927), 34:2 (954.5076) and 36:1 (m/z 982.5391) species. Data from stationary phase samples are similar. (B) Mass spectra obtained from integration of XIC peak region in (A) of Δ*wcaJ* (WT) and Δ*wcaJ* Δ*wecG* strains. WT peaks (black) are overlaid on top of Δ*wecG* peaks (red), illustrating unique DAG-PP-GlcNAc signals in the latter strain. (C) XICs of 34:1 DAG-PP-GlcNAc (m/z 956.5243) extracted from LC-MS analyses of IM lipids isolated from indicated Δ*wcaJ* strains. Inset, XICs of the same species from OM lipids.

**Fig. 8.**
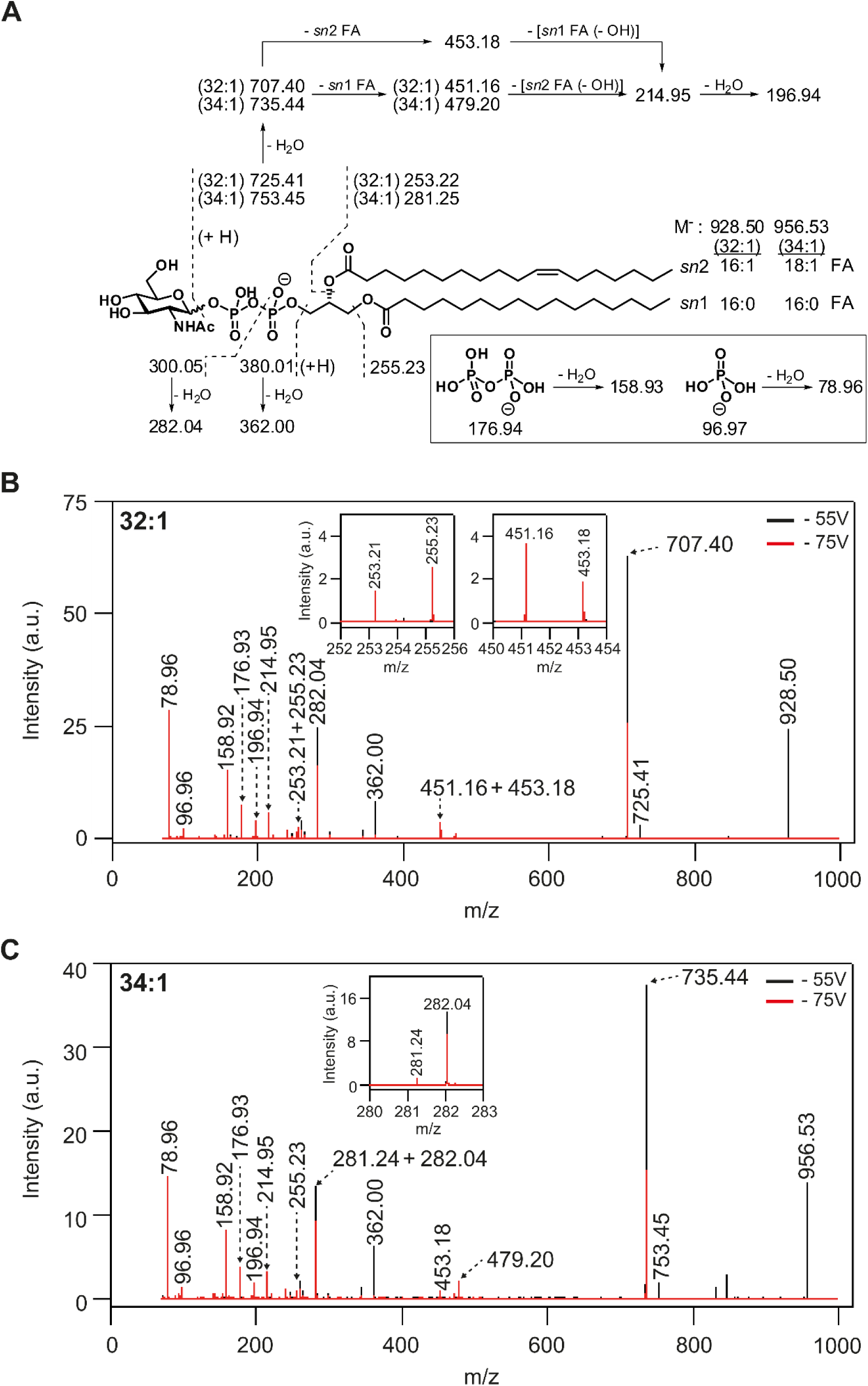
The chemical structures of DAG-PP-GlcNAc species are deduced from MS fragmentation analysis. (A) Proposed fragmentation pattern with mass assignments of 32:1 and 34:1 DAG-PP-GlcNAc species. (B and C) MS/MS spectra of (B) 32:1 and (C) 34:1 DAG-PP-GlcNAc species at indicated collision energies. Spectrum at high collision energy (−75 V, red) is overlaid on top of that at low collision energy (−55 V, black).

## Discussion

In this study, we have identified suppressor mutations that can rescue OM permeability defects in cells lacking the Tol-Pal complex, which are known to contain excess PLs in their OM (Shrivastava et al., 2017; Masilamani et al., 2018). Mutations in ECA biosynthesis restore OM barrier function, thus blocking entry of large antibiotics and reducing periplasmic leakiness, in *tol-pal* mutant strains (Figs. 1, 2); however, lipid homeostasis was not restored in these strains (Fig. 3). We have demonstrated that these rescue phenotypes are due to the accumulation of intermediate species along the ECA pathway (Figs. 4, 5). Interestingly, aside from the und-PP-linked ECA intermediates (Fig. 6), DAG-PP-linked species have also been observed (Figs. 7, 8). How the presence of these ECA-related species could result in the restoration of OM barrier function requires further investigation.

The role of ECA in the enterobacterial cell envelope is not known. Strains that do not make ECA are more sensitive to bile salts and produce more OMVs (Ramos-Morales et al., 2003; McMahon et al., 2012). Loss of cyclic ECA has also been reported to reverse OM barrier defects in strains lacking YhdP, a protein of unknown function (Mitchell et al., 2018). These observations suggest that ECA has roles related to the OM. However, we have shown that loss of any form of ECA (ECA_PG_/ECA_LPS_/ECA_CYC_) does not contribute to the restoration of OM barrier function in cells lacking the Tol-Pal complex; instead, the build-up of ECA biosynthetic intermediates is necessary for this rescue. Interestingly, accumulation of und-PP-linked intermediates (in the ECA, O-antigen, and colanic acid pathways) can lead to sequestration of und-P (Jorgenson et al., 2016; Jorgenson & Young, 2016), the common lipid carrier also used for peptidoglycan precursors. This can give rise to morphological changes due to a weakened cell wall (Figs. S9A, B). Overexpression of MurA to alleviate und-P sequestration effects reduced the extent of morphological defects but could not reverse the suppression of *tol-pal* phenotypes (Fig. S9). Intuitively, weakening the cell wall via und-P sequestration is unlikely to contribute to vancomycin and bacitracin resistance. We therefore believe that restoration of OM phenotypes by ECA intermediates in cells lacking the Tol-Pal complex is independent of und-P sequestration. Furthermore, the rescue mechanism does not appear to involve the major cell envelope stress responses, including Rcs, Cpx, and σ^E^ pathways (Figs. 2, S10).

One possible mechanism where accumulation of und-PP-linked ECA intermediates could affect the physical properties of the OM is via LPS modification. The structure of LPS could affect mechanical strength and barrier function of the OM (Rojas et al., 2018). In wild-type cells, the full length ECA polymer can be transferred onto LPS core oligosaccharides by WaaL to form ECA_LPS_ (Schmidt et al., 1976). However, since deletion of either *wecA* (does not restore OM barrier function) or other *wec* genes (restore OM barrier function) would result in the loss of full length ECA_LPS_, we ruled out the role of ECA_LPS_ in rescuing the OM permeability defect. It has been reported that the GlcNAc moiety from und-PP-GlcNAc (Lipid I^ECA^) can itself be transferred onto LPS in a WaaL-dependent manner (Feldman et al., 1999; Ruan et al., 2018). We reasoned it is likely that the incomplete sugar subunits on Lipid II/III^ECA^ could also be ligated onto LPS, therefore affecting the physical properties of the OM via LPS modification. However, we showed that deletion of *waaL* in strains that accumulate ECA intermediates did not reverse the suppression of OM permeability defects in the *tol-pal* mutant (Fig. S8). Therefore, the rescue of OM defects by ECA mutations in *tol-pal* strains is unlikely due to WaaL-dependent LPS modifications.

While und-PP-linked intermediates are expected to be accumulated in ECA mutants, we and others have also observed the corresponding DAG-PP-linked species (Rick et al., 1998). Specifically, DAG-PP-GlcNAc and DAG-PP-GlcNAc-ManNAc species have been detected in strains that accumulate und-PP-GlcNAc (Lipid I^ECA^) (Fig. 7) and und-PP-GlcNAc-ManNAc (Lipid II^ECA^) (Rick et al., 1998), respectively. Intriguingly, these unusual DAG-PP-linked species have been suggested to be precursors for the membrane protein integrase (MPIase) glycolipid, which in fact has a polymer of 9-11 trisaccharide units (identical to the ECA sugars) linked to the DAG-PP carrier (Nishiyama et al., 2012). It has been shown that MPIase is important for driving integration of membrane proteins into the IM, and also Sec-dependent translocation of periplasmic and OM proteins across the membrane (Nishiyama et al., 2012; Sawasato et al., 2019b). MPIase biosynthesis has been shown to require the activity of CdsA (Sawasato et al., 2019a), the CDP-diacylglycerol synthase critical for PL synthesis; however, despite the identical trisaccharide units, this process appears to be independent of the ECA pathway (Kamemoto et al., 2019). Consequently, it is not clear how and why DAG-PP-linked species, the presumed precursors to MPIase, would accumulate in ECA biosynthesis mutants. Regardless, we need to consider possible effects of these unusual lipids. We therefore speculate that these DAG-PP-linked species detected in our suppressor strains might somehow modulate the essential membrane protein integration or protein translocation functions of MPIase, possibly affecting a broad spectrum of pathways, and thus indirectly resulting in restored OM barrier function even in the event of lipid dyshomeostasis. The interesting connections between ECA, DAG-PP-linked species, and MPIase, remain to be clarified.

Cells lacking the Tol-Pal complex exhibit both cell division and OM stability/permeability defects. The cell division defect, which is more apparent under extreme osmolarity conditions, is thought to be a problem in OM invagination (Gerding et al., 2007) but recently shown to be also related to incomplete septal cell wall separation (Yakhnina & Bernhardt, 2020). The OM defects are largely independent of cell division (Yakhnina & Bernhardt, 2020), and likely due to lipid dyshomeostasis, where excess PLs accumulate in the OM (Shrivastava et al., 2017). While excess PLs in the OM necessarily affect lipid asymmetry (Shrivastava et al., 2017) and result in PL bilayer patches that can allow diffusion of hydrophobic molecules, it is unclear how that would increase permeation of large hydrophilic molecules, such as vancomycin, bacitracin, and RNase I, across the OM. One possibility is that having more PLs in the OM alters physical properties such as membrane tension and rigidity. These effects could in turn result in the appearance of transient “cracks” in the bilayer, which have been suggested to allow the passage of large hydrophilic compounds (Nikaido, 2005; Ruiz et al., 2006). Vancomycin can also cross the OM via the lumen of large porins, if unplugged (Krishnamoorthy et al., 2016). Therefore, another possibility is that the changes in lipid composition in the OM of cells lacking the Tol-Pal complex may affect the structure and function of specific large OM porins, causing them to become leaky. How the accumulation of und-PP-linked ECA intermediates and/or DAG-PP-linked species restores OM barrier function against large hydrophilic molecules remains unclear; however, these species might affect other yet-to-be-identified pathways (e.g. MPIase function), possibly indirectly modifying OM physical properties or porin function. The unappreciated roles of ECA-related species in OM biology should be thoroughly investigated.

### Experimental Procedures

#### Strains and growth conditions

All the strains used in this study are listed in Table S2. *Escherichia coli* strain MC4100 [*F*^*-*^ *araD139* Δ*(argF-lac) U169 rpsL150 relA1 flbB5301 ptsF25 deoC1 ptsF25 thi*] (Casadaban, 1976) was used as the wild-type (WT) strain for most of the experiments. NR754, an *araD*^+^ revertant of MC4100 (Ruiz et al., 2008), was used as the WT strain for experiments involving depletion of *wzxE* or *wzyE* from the arabinose-inducible promoter (P_BAD_). Gene deletion mutants were constructed using recombineering (Baba et al., 2006; Datsenko et al., 2000) or obtained from the Keio collection (Baba et al, 2006). Whenever needed, the antibiotic resistance cassettes were flipped out as described (Baba et al., 2006; Datsenko et al., 2000). Gene deletion cassettes were transduced into relevant genetic background strains via P1 transduction (Silhavy et al., 1984). The unmarked and chromosomal *wecC*\* allele was constructed using a negative selection technique (Khetrapal et al., 2015). Luria-Bertani (LB) broth (1% tryptone and 0.5% yeast extract, supplemented with 1% NaCl) and agar were prepared as previously described (Silhavy et al., 1984). When appropriate, kanamycin (kan; 25 µg ml^−1^), chloramphenicol (cm; 30 µg ml^−1^), ampicillin (amp; 200 µg ml^−1^) and spectinomycin (spec; 50 µg ml^−1^) were added.

#### Plasmid construction

Plasmids used in this study are listed in Table S3. Desired genes were amplified from MC4100 chromosomal DNA using the indicated primers (sequences in Table S4). Amplified products were digested with indicated restriction enzymes (New England Biolabs), which were also used to digest the carrying vector. After ligation, recombinant plasmids were transformed into competent NovaBlue (Novagen) cells and selected on LB plates containing appropriate antibiotics. DNA sequencing (Axil Scientific, Singapore) was used to verify the sequence of the cloned gene.

#### Generation of suppressor mutations and genome sequencing

To isolate spontaneous suppressor mutants, 10^9^ BW25113 Δ*tol-pal*^*#*^ cells were plated on LB agar plate supplemented with vancomycin (250 μg/ml) and incubated at 37°C for 48 h. Individual colonies were picked and restreaked on similar plates to verify their vancomycin resistance properties. 36 separate strains were isolated. To identify the genetic location of mutations, whole genome sequencing was performed. Purified genomic DNA was sheared to approximately 300 bp using a focused ultrasonicator (Covaris). A sequencing library was prepared using the TruSeq DNA PCR Free Kit (Illumina) according to the manufacturer’s instructions. This was sequenced using a HiSeq 4000 with 2×151 bp reads. Raw FASTQ files were mapped to the *E. coli* W3110 genome sequence (NC_007779.1) using bwa (version 0.7.10) (Li & Durbin, 2009); indel realignment and SNP (single nucleotide polymorphism) calling was performed using Lofreq* (version 2.1.2) with default parameters (Wilm et al., 2012). Resulting variants were assigned to associated genes and amino acid changes using the Genbank Refseq W3110 annotation.

#### Antibiotic sensitivity assay

Sensitivity against different antibiotics was judged by efficiency of plating (EOP) analyses on LB agar plates containing indicated concentrations of drugs. Vancomycin (V1130-1G) and bacitracin (11702-5G) are acquired from Sigma Aldrich. Briefly, 5-ml cultures were grown (inoculated with overnight cultures at 1:100 dilution) in LB broth at 37°C until OD_600_ reached ∼0.6. Cells were normalized according to OD_600_, first diluted to OD_600_ = 0.1 (≈10^8^ cells), and then serially diluted in LB with six 10-fold dilutions using 96-well microtiter plates (Corning). Two microliters of the diluted cultures were manually spotted onto the plates and incubated overnight at 37°C. All results shown are representative of at least three independent replicates.

#### RNase I leakage assay

Measurement of RNase I leakiness was performed using a plate assay as described before (Lazzaroni & Portalier, 1979). Briefly, 5-ml cultures were grown (inoculated with overnight cultures at 1:100 dilution) in LB broth at 37°C until OD_600_ reached ∼0.6. Cells were normalized to OD_600_ = 0.001, and 2 μl (≈2,000 cells) were manually spotted onto LB agar plates containing 1.9 mg/ml yeast RNA extract (Sigma). The plates were incubated overnight at 37°C. To precipitate and visualize RNA, the plates were overlaid with cold (12.5% v/v) trichloroacetic acid. Size of the halo were defined as the distance between the edge of the macrocolony and the edge of the halo.

#### Microscopy

5-ml cultures were grown (inoculated with overnight cultures at 1:2000 dilution) in LB broth at 37°C until OD_600_ reached ∼0.5 – 0.6. For FM1-43 (Invitrogen) labelling, 1 μg/ml final concentration of the probe was added to 100 μl of bacterial culture before imaging. For vancomycin-BODIPY FL (Invitrogen) labelling, cell cultures were incubated with 10 μg/ml final concentration of the probe for 15 minutes and washed one time with 1 ml fresh LB before resuspended into 200 μl LB. For all experiments, 5 μl of final cell culture were spotted onto freshly prepared 1% LB agarose pads. Images were acquired using a Zeiss LSM710 confocal microscope at 100x magnification. Cells lengths and widths were measured with Image J (Schneider et al., 2012) and result plotted using the *ggplot2* package in R.

#### RT-PCR

Strains with empty vector or the *uppS/murA* overexpression plasmid were grown in the presence of 25 μM IPTG to mid-log phase. For each strain, total RNA was isolated using RNeasy Mini Kit (Qiagen) and corresponding complementary DNA library was prepared using QuantiTect Reverse Transcription Kit (Qiagen). cDNA sample for each strain was normalized and used as template for PCR amplification using primers specific to *uppS, murA*, and *gyrA* (housekeeping control) for different number of cycles (26, 36, and 23 respectively) to prevent saturation. Following gel electrophoresis, band intensity of the resulting *uppS* and *murA* PCR products was normalized to the corresponding *gyrA* PCR product to obtain the relative gene expression level.

#### σ^E^ reporter assay

Strains with plasmid expressing *rpoH*P3::*gfp* or promoterless *gfp* were grown to mid-logarithmic phase (OD_600_ ≈ 0.4-0.6), then normalized and re-suspended in 150 mM NaCl. Fluorescence level (Ex: 485 nm and Em: 535 nm) of equal amount of cells were measured for each strain using the Victor X4 plate reader (Perkin Elmer). Relative levels of σ^E^ activation correlates with the expression of GFP due to specific activation of *rpoH*P3 promoter and were determined by normalizing the fluorescence level in strains with *rpoH*P3::*gfp* to the basal fluorescence level in strains expressing *gfp* without promoter. Data from three independent experiments were collected and normalized to the σ^E^ activation level in WT.

#### Lipid extraction and liquid chromatography-mass spectrometry (LC-MS) analysis

To prepare the lipid extracts, a modified Bligh and Dyer method was used as described previously (Guan & Maser, 2017). Briefly, bacterial pellets were resuspended in PBS, and chloroform:methanol (1:2, v/v) was added. The mixture was vortexed thoroughly before incubation with shaking at 1,000 rpm, 4°C. Subsequently, water and chloroform were added to each sample to generate a two-phase Bligh-Dyer mixture. The two phases were separated via centrifugation and the lower organic phase was collected in a new tube. The aqueous phase was re-extracted twice with chloroform, and all the organic extracts pooled and dried using a Centrivap and stored at −80°C until use.

The lipid samples were reconstituted in chloroform:methanol (1:1, v/v) and analyzed using a high performance chromatography system (1260 Agilent Infinity Quaternary Pump) coupled to an SCIEX QTOF 6600 mass spectrometer in negative electrospray ionization mode. Mass calibration is performed every 5 h, using the automated calibration solution (SCIEX, Canada). For lipid separation, normal phase chromatography was performed as previously described (Guan & Maser, 2017). For characterization of the DAG-PP species using tandem mass spectrometry, multiple collision energies ranging from −55 V to −85 V were used. MS and MS/MS spectra obtained were visualized using Peak View (SCIEX) and graphical representations of the selected peaks of interests were plotted using sigmaplot v10.0.

#### Membrane lipid composition analyses

Experiments to determine steady-state radiolabeled PL distributions in IMs/OMs and PL/LPS ratios in OMs were adapted from methods previously described with some changes (Shrivastava et al., 2017). Briefly, 10 ml cultures were grown in LB broth (inoculated from overnight culture to a starting OD_600_ of 0.05) containing [^32^P]-disodium phosphate (final 1 µCi ml^−1^; Perkin Elmer product no. NEX011001MC) until mid-log phase (OD_600_ ∼0.5–0.7). Cells were harvested by centrifugation at 4700 × *g* for 10 min, re-suspended in 5 ml of TBS (20 mM pH 8.0 Tris-HCl, 150 mM NaCl), and centrifuged again as above. Resulting cell pellets were re-suspended in 5 ml of 20% sucrose in 10mM Tris-HCl pH 8.0 (w/w) containing 1 mM PMSF and 50 μg ml^−1^ DNase I), and lysed by a single passage through a high pressure French press (French Press G-M, Glen Mills) homogenizer at 8000 psi. Unbroken cells were removed by centrifugation at 4700 × *g* for 10 min. The cell lysate was collected, and 5.5 ml of cell lysate was layered on top of a two-step sucrose gradient consisting of 40% sucrose solution (5 ml) layered on top of 65% sucrose solution (1.5 ml) at the bottom of the tube. Samples were centrifuged at 36 000 rpm for 16 h in a Beckman SW41 rotor in an ultracentrifuge (Model XL-90, Beckman). 0.8-ml fractions (usually 15 fractions) were manually collected from the top of each tube. IM and OM fractions were pooled and diluted with TBS before centrifugation at 36 000 rpm for one hour in the Beckman SW41 rotor to concentrate the membrane. The resulting membrane pellets (not visible) were resuspended in 320 µl TBS for subsequent extraction of PLs (IM) or PLs and LPS (OM). Protocol for the extraction of PLs and LPS had been described previously (Shrivastava et al., 2017). Dried PLs were resuspended in 50 µl of a mixture of chloroform:methanol (2:1); while dried LPS pellets were resuspended in 50 µl 1% SDS. Equal volumes of PL or LPS solutions were mixed with 2 ml of Ultima Gold scintillation fluid (Perkin Elmer, Singapore) and [^32^P]-counts were measured using scintillation counting (MicroBeta^2®^, Perkin-Elmer). For each strain, scintillation counts of OM PLs were divided by the total counts of IM PLs and OM PLs to obtain the percentage of OM PLs distribution; while scintillation counts of OM PLs were divided by the counts of OM LPS to obtain the PL/LPS ratio.

#### Quantification of OM vesiculation

For each strain, 10-ml cells were grown at 37°C in LB broth (inoculated from an overnight culture at 1:100 dilution) containing [1-^14^C]-acetate (final 0.2 µCi ml^−1^; Perkin Elmer product no. NEC084A001MC) until OD600 reached ∼0.7. At this OD, cultures were harvested to obtain the cell pellets, and supernatants containing OM vesicles. Cell pellets were washed twice with 10 mM Tris-HCl pH 8.0 and finally suspended in the same buffer (0.2 ml). To obtain OM vesicles, supernatants were filtered through 0.45 μm filters followed by ultracentrifugation in a SW41.Ti rotor at 39,000 rpm for 1 h. Finally, the OM vesicles in the resulting pellets were washed and re-suspended in 0.2 ml 10 mM Tris-HCl pH 8.0 buffer. Radioactive counts in cell pellets and OM vesicles were measured after mixing with 2 ml of Ultima Gold scintillation fluid (Perkin Elmer, Singapore). Radioactivity ([^14^C]-count) was measured on a scintillation counter (MicroBeta^2®^, Perkin-Elmer)

#### SDS-PAGE and immunoblotting

SDS-PAGE was performed according to Laemmli using the 12% or 15% Tris.HCl gels (Laemmli, 1970). For ECA, cell samples were treated with proteinase K (0.25 mg/ml) at 55°C for 1 h before loading and resolving on 10% Tricine SDS-PAGE gels. Immunoblotting was performed by transferring from the gels onto polyvinylidene fluoride (PVDF) membranes (Immun-Blot® 0.2 μm, Bio-Rad) using the semi-dry electroblotting system (Trans-Blot® TurboTM Transfer System, Bio-Rad). Membranes were blocked using 1X casein blocking buffer (Sigma). Rabbit polyclonal α-ECA_CYC_ antisera (generous gift from Jolanta Lukasiewicz), which reacts with all forms of ECA (ECA_CYC_, ECA_LPS_, ECA_PG_), was used at 1:800 dilution (Gozdzlewicz et al., 2014). α-rabbit IgG secondary antibody conjugated to HRP (from donkey) was purchased from GE Healthcare and used at 1:5,000 dilutions. Luminata Forte Western HRP Substrate (Merck Milipore) was used to develop the membranes and chemiluminescent signals were visualized by G:BOX Chemi XT 4 (Genesys version1.3.4.0, Syngene).

## Acknowledgements

We thank Majid Eshaghi (Genome Institute of Singapore, GIS) for providing σ^E^ reporter plasmids and Kevin Young (University of Arkansas for Medical Sciences) for the generous gifts of the Δ*wcaJ* strain and the *murA* and *uppS* overexpression plasmids. We also thank Kevin Young for critical comments. We are grateful to Jolanta Lukasiewicz (Polish Academy of Sciences) for the α-ECA antibody. WGS work was partially supported by the GIS, and the Singapore Ministry of Health National Medical Research Council (NMRC/CIRG/1357/2013) to S.L.C.. Lipid MS analysis was supported by the Nanyang Assistant Professorship to X.L.G.. All other work were supported by the Singapore Ministry of Education Academic Research Fund Tier 1 Grant, and the Singapore Ministry of Health National Medical Research Council under its Cooperative Basic Research Grant (NMRC/CBRG/0072/2014) to S.-S.C.. The authors declare no conflict of interest.

## Author contributions

X.E.J., W.B.T., and R.S. performed most of the experiments described in this work; S.L.C. analyzed the whole genome sequencing data of suppressor mutants; D.C.C.S. and X.L.G. performed the MS experiments; S.-S.C. directed and supervised the work; X.E.J., R.S., W.B.T., and S.-S.C. analyzed the data and wrote the paper; all authors provided critical feedback of the manuscript.

## Data availability statement

The data that supports the findings of this study are available in the supplementary material of this article and from the corresponding author upon reasonable request.

