## Supplementary figures and tables for "Mutations in enterobacterial common antigen biosynthesis restore outer membrane barrier function in *Escherichia coli tol-pal* mutants"

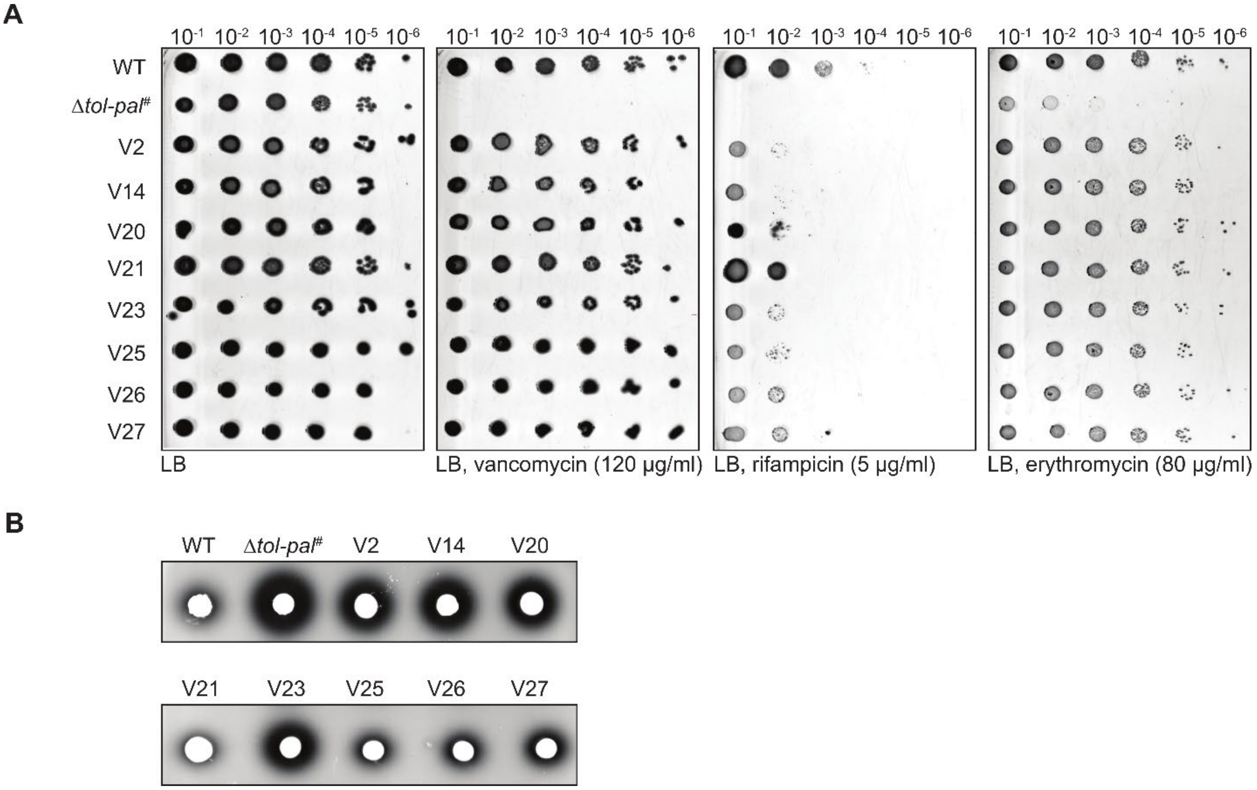


**Fig. S1** Suppressor mutants of cells lacking the Tol-Pal complex have partially restored OM barrier function. (A) Antibiotic sensitivities of WT, Δ*tol-pal*#, and the indicated suppressor mutants based on EOP on LB agar plates supplemented with vancomycin (120 μg/ml), rifampicin (5 μg/ml), or erythromycin (80 μg/ml). (B) RNase I leakage in the same strains as (A), as judged by RNA degradation (halo formation) around cells spotted on LB agar plates containing yeast RNA, subsequently precipitated with trichloroacetic acid.

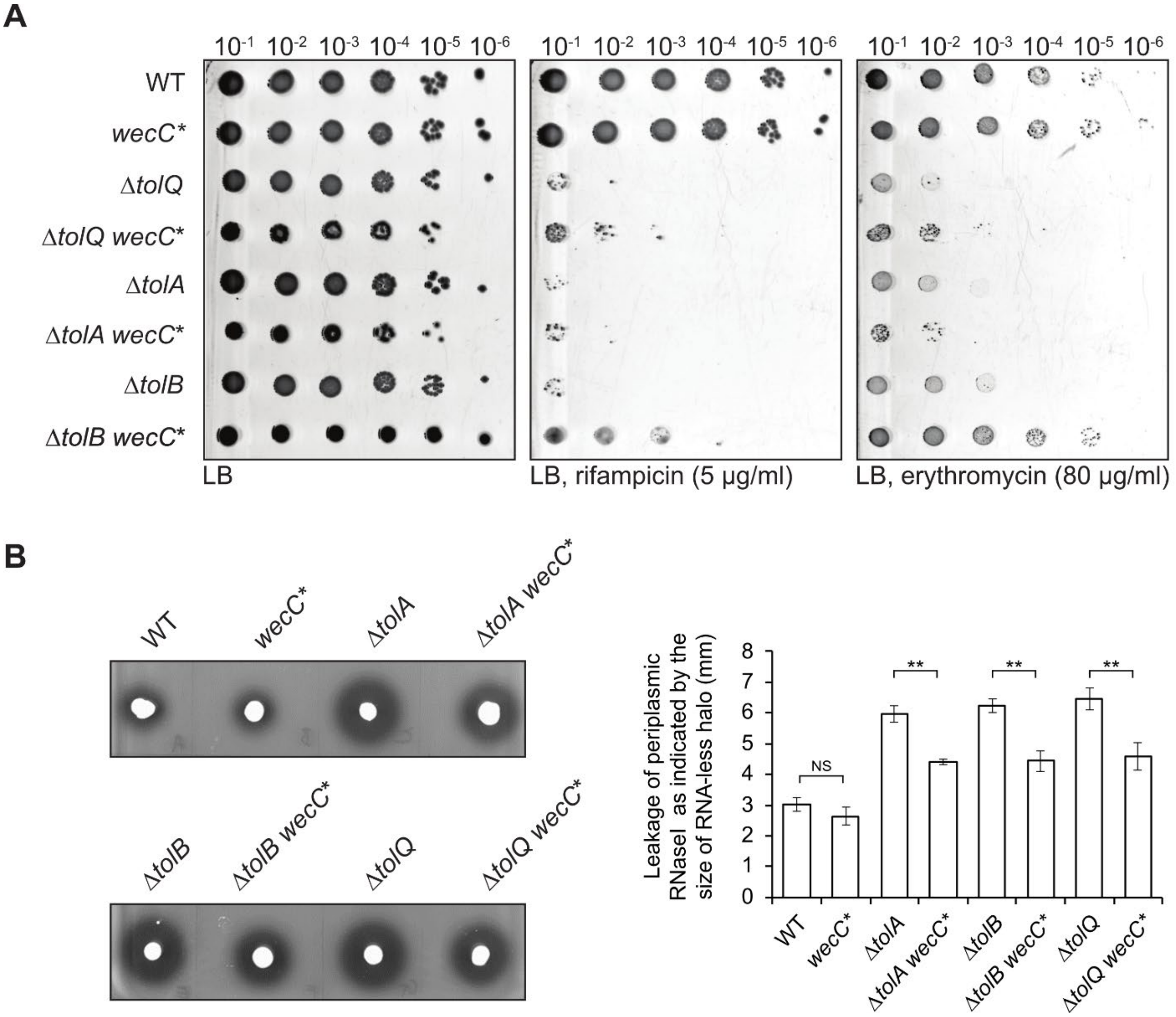


**Fig. S2** The *wecC** mutation partially reduces sensitivity to rifampicin and erythromycin and RNase I leakage in cells lacking the Tol-Pal complex. (A) Antibiotic sensitivities of the indicated strains, based on EOP on LB agar plates supplemented with rifampicin (5 µg/ml), or erythromycin (80 µg/ml). (B) RNase I leakage of the same strains, as judged by RNA degradation (halo formation) around cells spotted on LB agar plates containing yeast RNA, subsequently precipitated with trichloroacetic acid. Quantification of the distances between the edges of the macrocolony and the halo ($n = 4$) is shown on the right; error bars represent standard deviation. Student's t-tests: *, $p < 0.05$; **, $p < 0.005$.

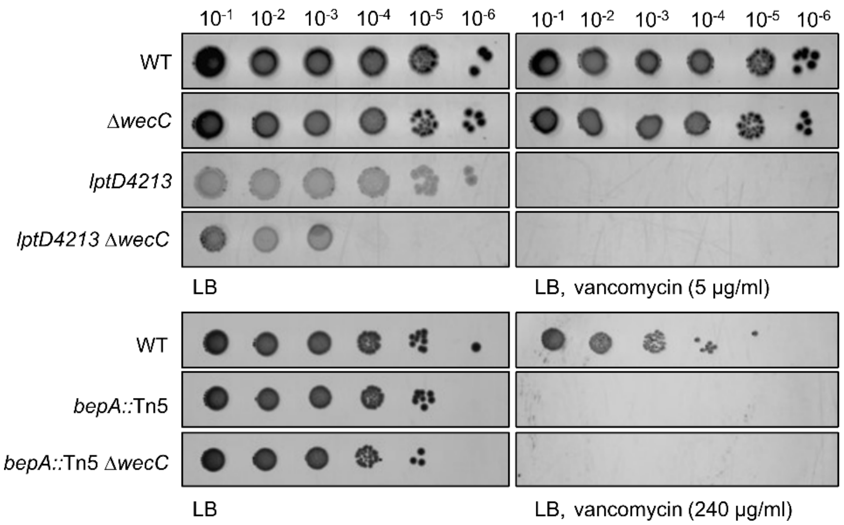


**Fig. S3** Δ*wecC* does not restore OM permeability defects caused by *lptD4213* and *bepA*::Tn5 mutations, as indicated by the EOP of the respective strains on LB agar plates containing vancomycin at the indicated concentrations.

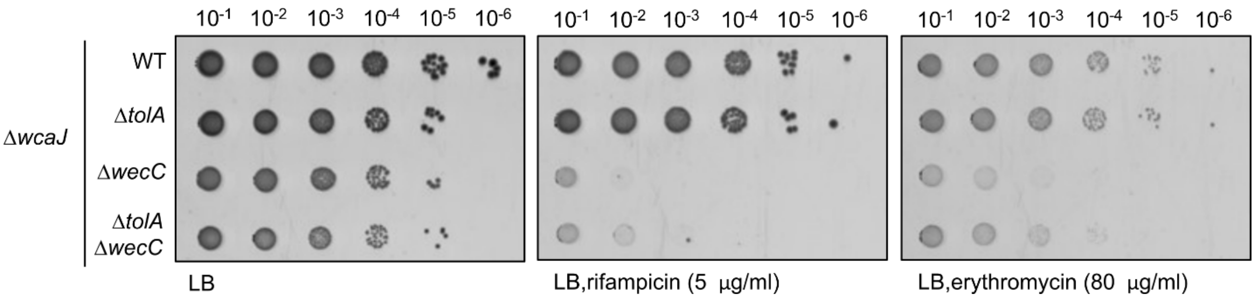


**Fig. S4** Δ*wecC* partially rescues sensitivity to rifampicin and erythromycin in cells lacking the Tol-Pal complex in the Δ*wcaJ* background. Antibiotic sensitivities of the indicated Δ*wcaJ* strains based on EOP on LB agar plates supplemented with rifampicin (5 μg/ml), or erythromycin (80 μg/ml).

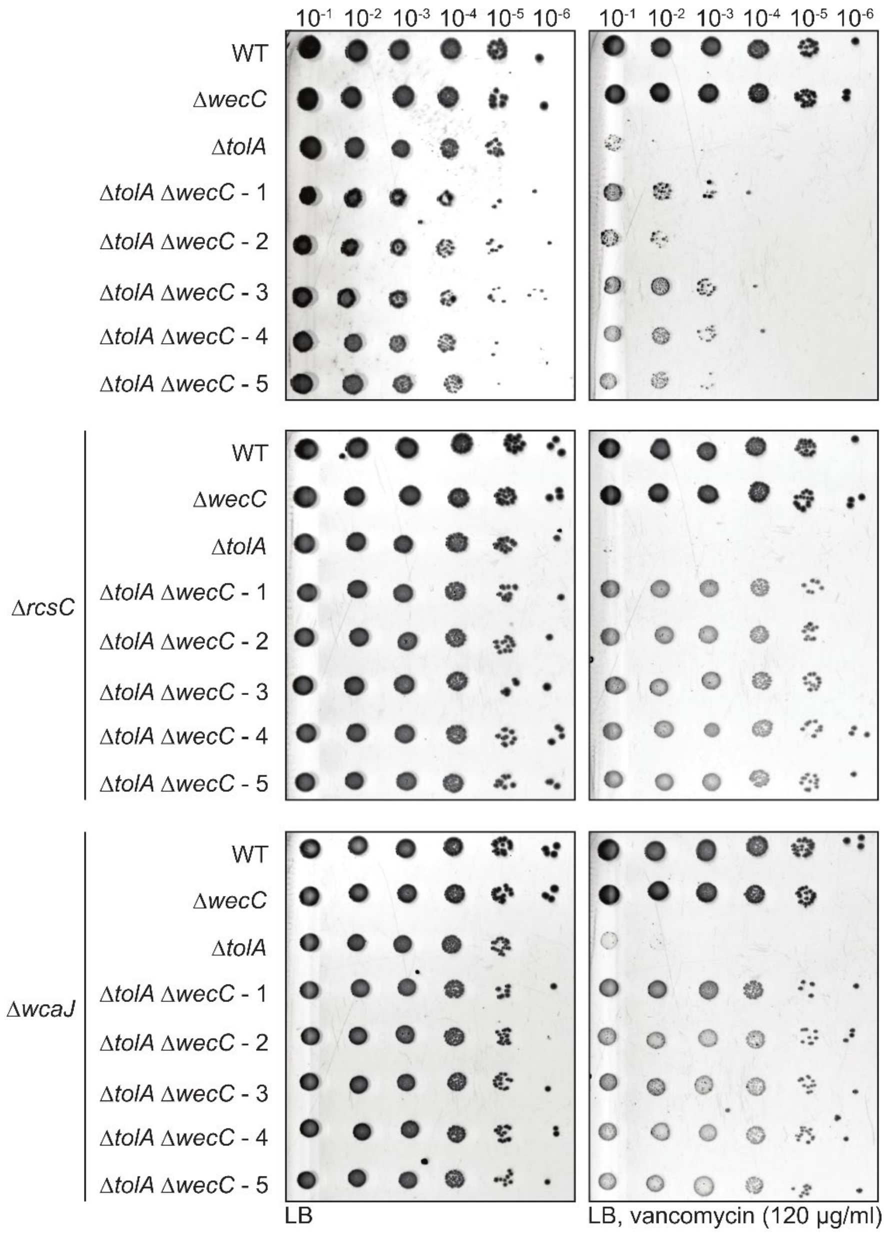


**Fig. S5** Removing RcsC or WcaJ improves suppression of vancomycin sensitivity by the Δ*wecC* mutation in the Δ*tolA* strain. Vancomycin sensitivity of WT, Δ*wecC*, Δ*tolA*, and distinct Δ*tolA* Δ*wecC* strains (1 – 5) in the WT (*top panel*), Δ*rcsC* (*middle panel*) or Δ*wcaJ* (*bottom panel*) background, based on EOP on LB agar plates supplemented with vancomycin (120 μg/ml).

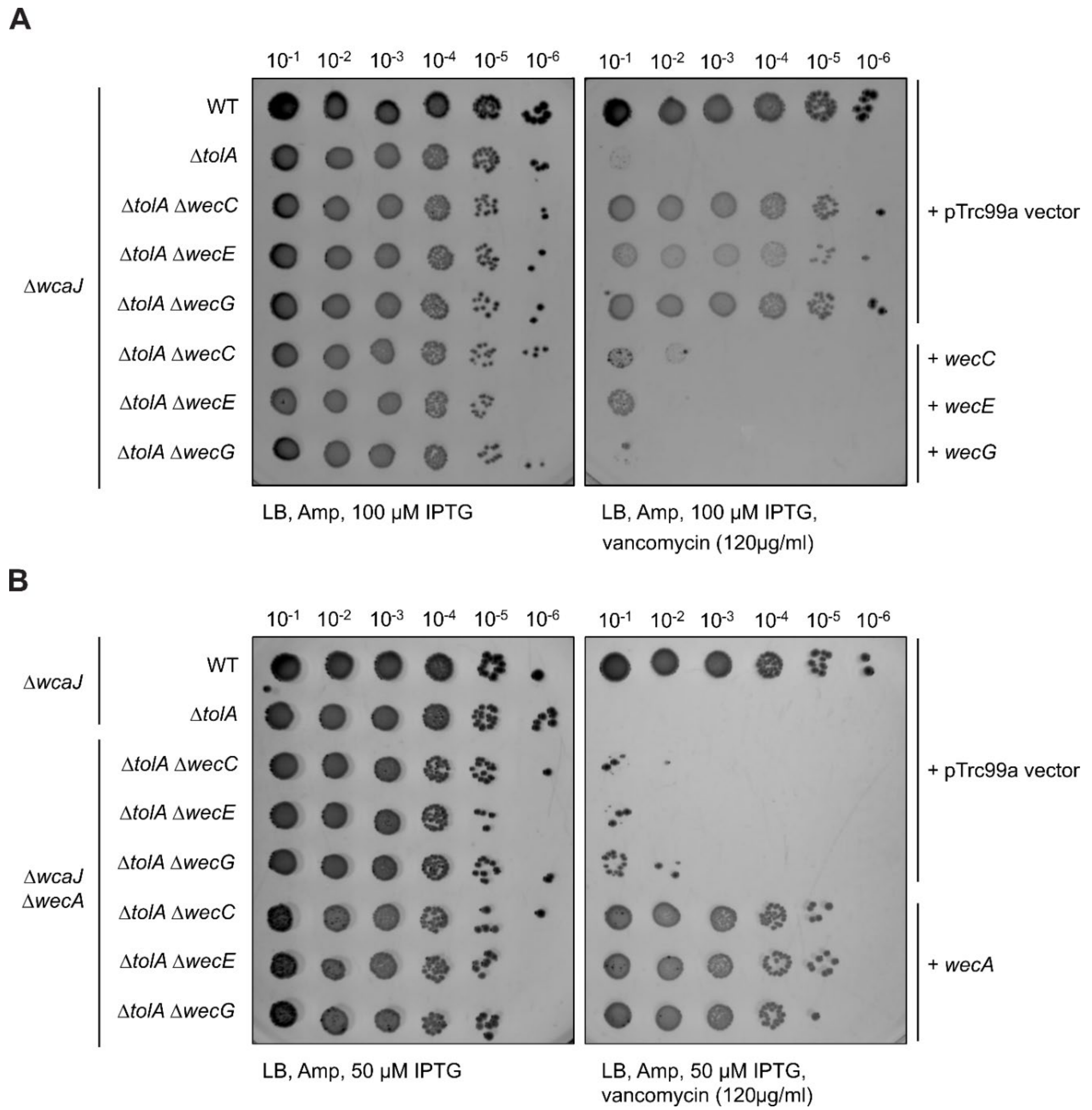


**Fig. S6** Expressing *wecC*/*E*/*G* or *wecA in trans* reverses the effects of Δ*wecC*/*E*/*G* and Δ*wecA* on the vancomycin sensitivity of the Δ*wcaJ* Δ*tolA* strains. (A) Vancomycin sensitivity of the Δ*wcaJ* Δ*tolA* Δ*wecC*/*E*/*G* strains harbouring empty vector or pTrc99a-*wecC/E/G*, based on EOP on LB agar plates supplemented with ampicillin, IPTG (100 μM), and vancomycin (120 μg/ml). (B) Vancomycin sensitivity of the Δ*wcaJ* Δ*tolA* Δ*wecA* Δ*wecC*/*E*/*G* strains harbouring empty vector or pTrc99a-*wecA*, based on EOP on LB agar plates supplemented with ampicillin, IPTG (50 μM), and vancomycin (120 μg/ml).

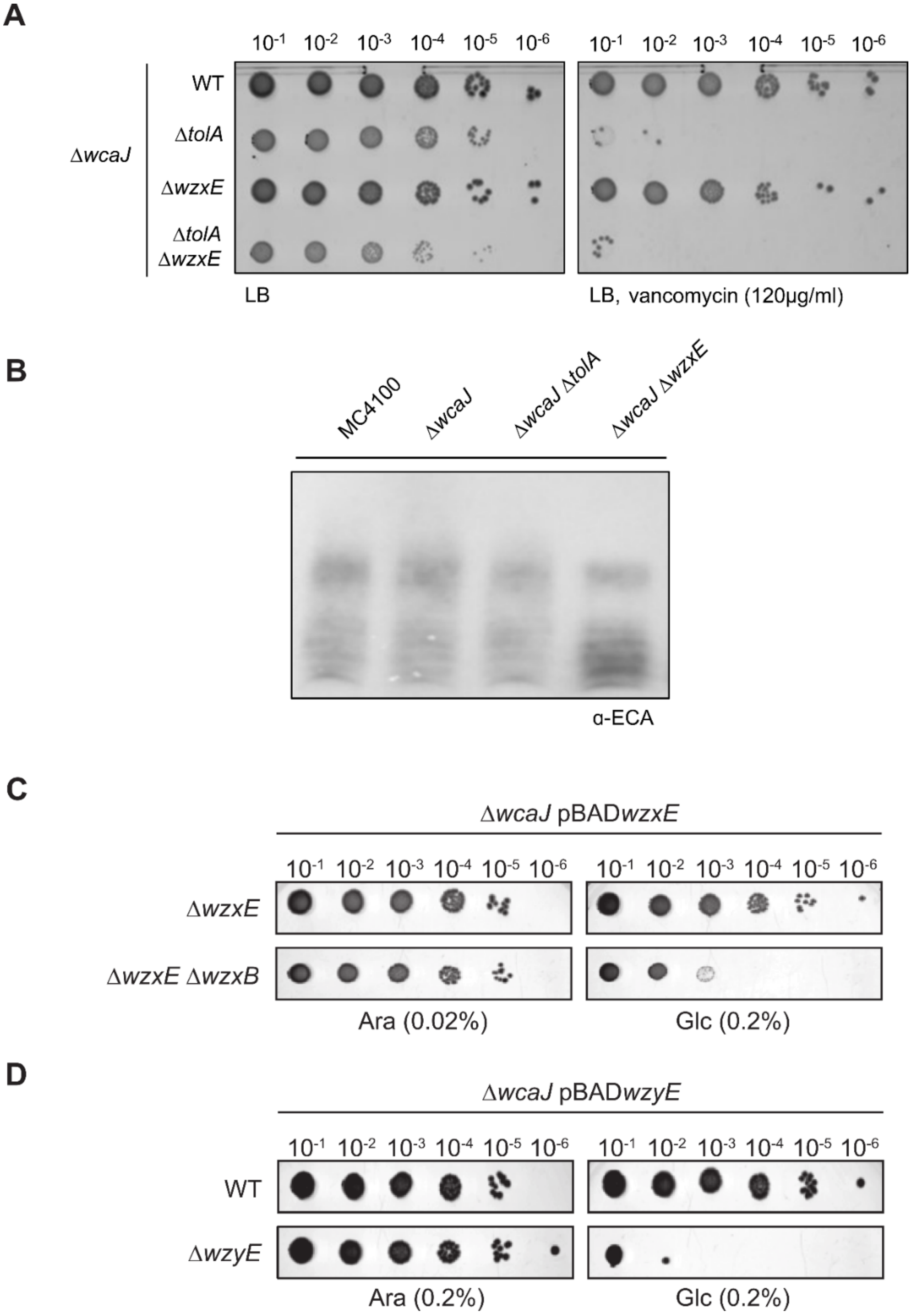


**Fig. S7** Depletion of *wzxE* or *wzyE*, leading to accumulation of Lipid III$^{ECA}$, is lethal. (A) Δ*wzxE* does not confer vancomycin resistance to Δ*wcaJ* Δ*tolA* strain, based on EOP on LB agar plates supplemented with ampicillin and vancomycin (120 μg/ml). (B) Δ*wzxE* does not abolish production of ECA, as judged by immunoblot analysis using α-ECA antibody. Growth defects of (C) *wzxE* depletion in either Δ*wcaJ* Δ*wzxE* or Δ*wcaJ* Δ*wzxE* Δ*wzxB* strains, or (D) *wzyE* depletion in either Δ*wcaJ* (WT) or Δ*wcaJ* Δ*wzyE* strains, based on EOP on LB agar plates supplemented with indicated concentrations of arabinose or glucose. *wzxE* and *wzyE* are expressed under the control of the arabinose-inducible promoter in the pBAD33 plasmid.

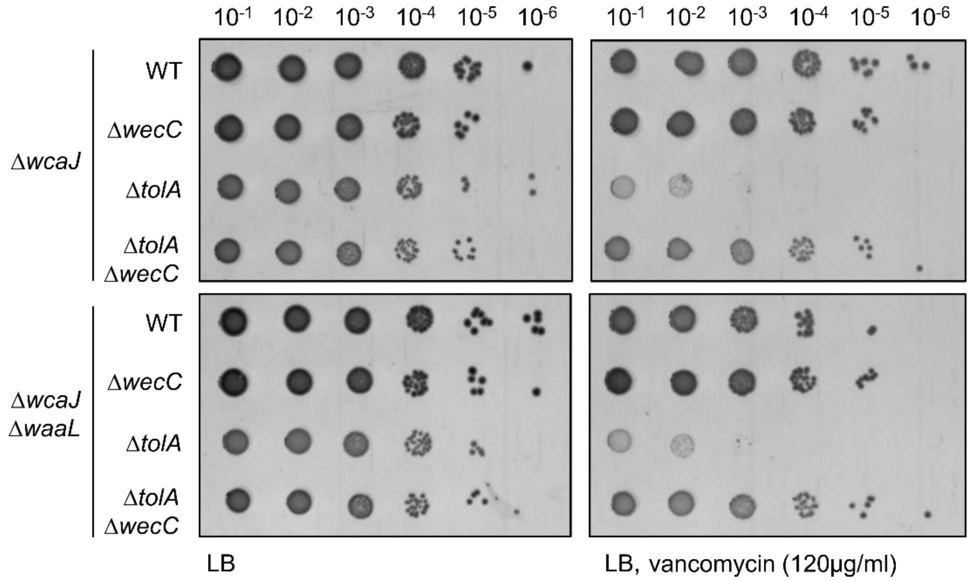


**Fig. S8** Restoration of the OM barrier function by Δ*wecC* is not dependent on WaaL. Deleting *waaL* in the Δ*wcaJ* Δ*tolA* Δ*wecC* strain does not revert the vancomycin resistance, based on EOP on LB agar plates supplemented with vancomycin (120 μg/ml).

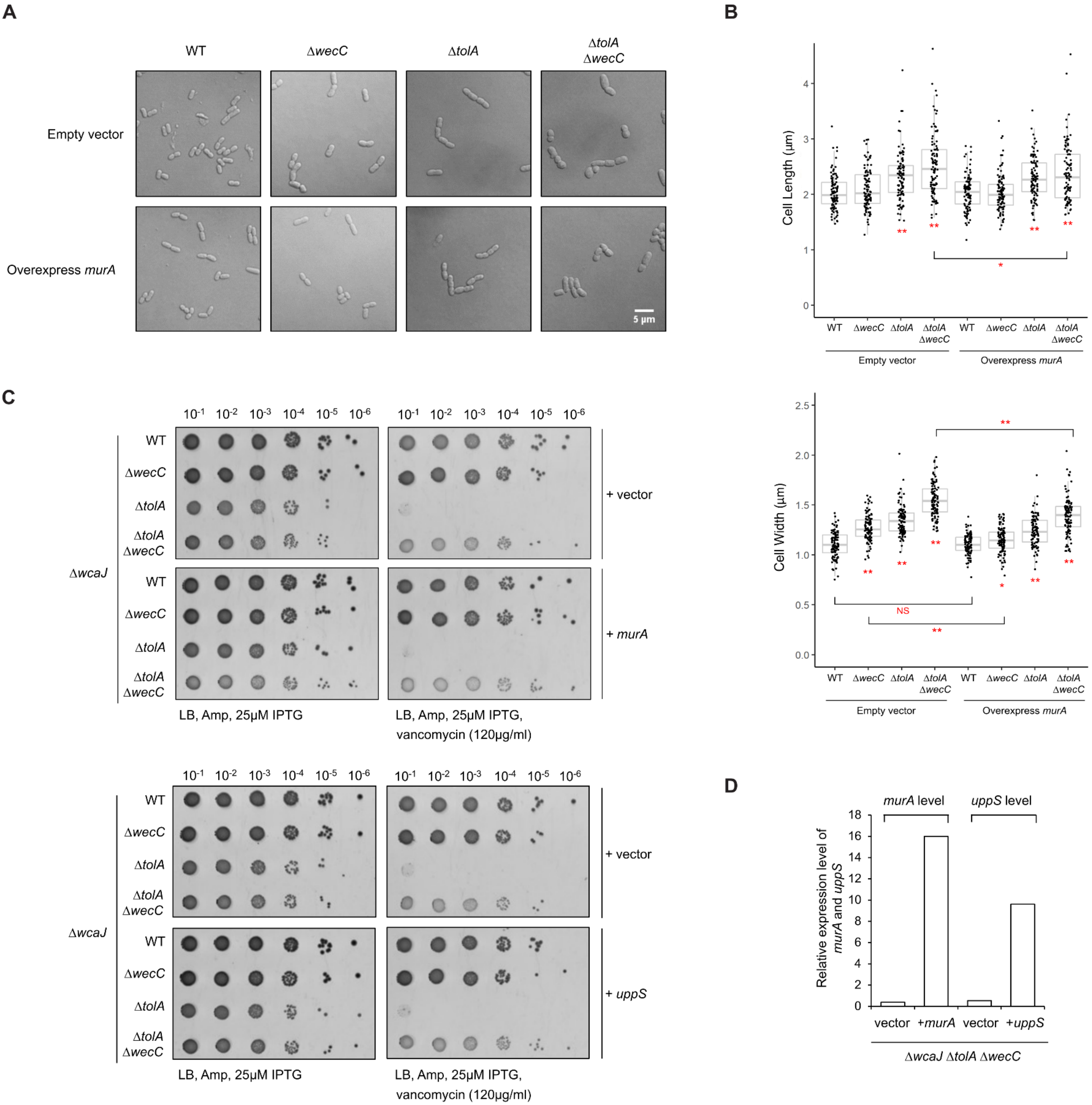
A
WT
ΔwecC
ΔtolA
ΔtolA ΔwecC
Empty vector
Overexpress murA
5 μm
B
Cell Length (μm)
Cell Width (μm)
WT ΔwecC ΔtolA ΔtolA ΔwecC WT ΔwecC ΔtolA ΔtolA ΔwecC
Empty vector
Overexpress murA
NS
C
10-1 10-2 10-3 10-4 10-5 10-6
ΔwcaJ
WT
ΔwecC
ΔtolA
ΔtolA ΔwecC
+ vector
+ murA
+ uppS
LB, Amp, 25μM IPTG
LB, Amp, 25μM IPTG, vancomycin (120μg/ml)
D
murA level
uppS level
Relative expression level of murA and uppS
vector
+murA
+uppS
ΔwcaJ ΔtolA ΔwecC

**Fig. S9** ECA intermediate-dependent restoration of vancomycin resistance in cells lacking TolA is not due to und-P sequestration. (A) Cell morphology of indicated Δ*wcaJ* strains visualized using DIC confocal microscopy. (B) Cell length and width of 100 cells from each sample in (A) were measured and plotted. Scale bar, 5 µm. Statistical analysis done using Wilcoxon ranked sum test; ∗, $p < 0.05$; ∗∗, $p < 0.005$; NS, not significant. Unless otherwise indicated, the *p*-value is for comparison against WT strain carrying the same vector. (C) Vancomycin sensitivity of indicated Δ*wcaJ* strains harboring the empty vector pDSW204, pDSW204*murA*, or pDSW204*uppS* based on EOP on LB agar plates containing vancomycin (120 µg/ml). 25 µM IPTG was added to induce overexpression of *murA* and *uppS*. (D) Relative transcript levels of *murA* or *uppS* (RT-PCR bands normalized against that of *gyrA*) in the Δ*wcaJ* Δ*tolA* Δ*wecC* strains with or without *murA* or *uppS* overexpression.

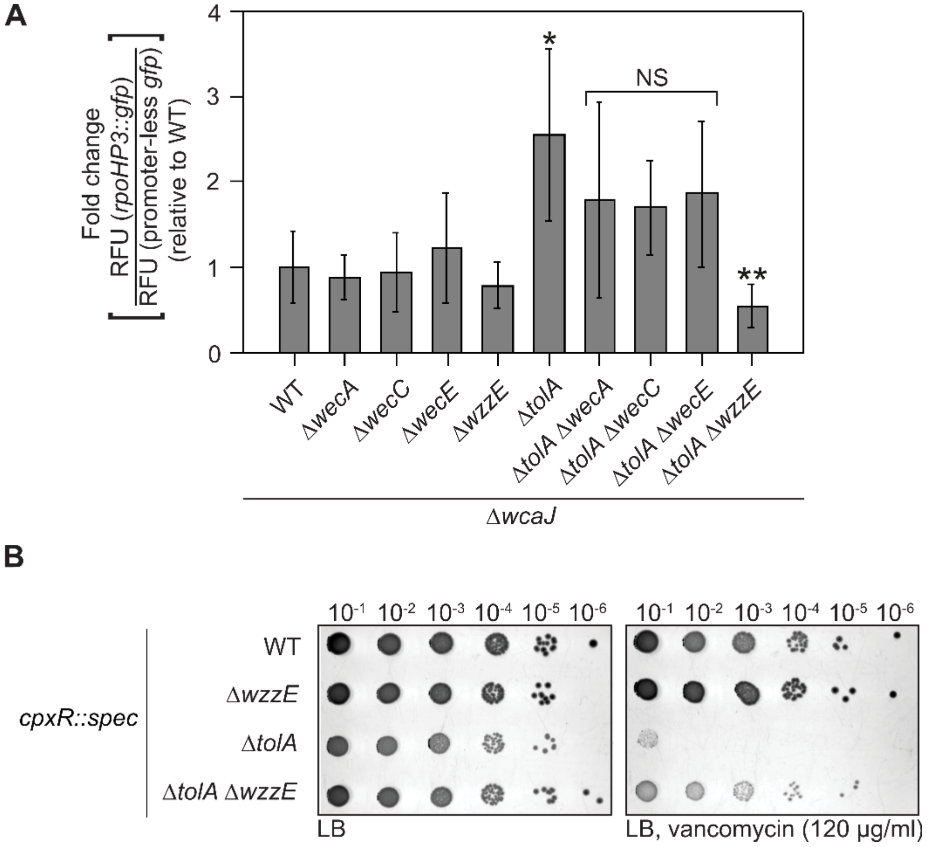


**Fig. S10** ECA intermediate-dependent restoration of vancomycin resistance in cells lacking TolA is independent of $\sigma^E$ and Cpx stress responses. (A) Relative levels of $\sigma^E$ activation in indicated Δ*wcaJ* strains as judged by expression of *rpoH*P3::*gfp* normalized to that of promoter-less *gfp*. Error bars represent standard deviations calculated from biological triplicates. Student's t-tests: **p* < 0.05 (as compared to WT); ***p* < 0.005 (as compared to Δ*tolA*); NS, not significant. (B) Vancomycin sensitivity of indicated Δ*wcaJ* strains in a *cpxR* background, based on EOP on LB agar plates containing vancomycin (120 μg/ml).

**Table S1 List of mutations in suppressor strains**

| Strain name | Mutated gene | Gene length (bp) | Mutation in DNA sequence | Change in protein |
|---|---|---|---|---|
| V2 | *wecB* | 1131 | $A_{743}C$ | N248T |
| | *pgm* | 1641 | $A_{352}C$ | T118P |
| V14 | *wecC* | 1263 | $C_{783}CCCTGGC$ | G261GPG |
| | *pgm* | 1641 | $C_{1532}T$ | T511M |
| | *mdoH* | 2544 | $T_{1381}TG$ | frame-shift |
| | *rpoS* | 993 | $A_{358}AT$ | frame-shift |
| V20 | *wecC* | 1263 | $A_{659}C$ | D220A |
| | *mdoH* | 2544 | $G_{176}A$ | W59X |
| | *araD* | 696 | $A_{23}G$ | Q8R |
| V21 | *wecF* | 1080 | $T_{239}TA$ | frame-shift |
| | *mdoH* | 2543 | $G_{176}A$ | W59X |
| | *prc* | 2049 | $CG_{812}C$ | frame-shift |
| | *nlpD* | 1140 | $C_{1132}CTGCGTTATTTGCCGC$ | Q377LRYLPQ |
| V23 | *wecC* | 1263 | $G_{775}T$ | G259C |
| | *mdoH* | 2544 | $G_{1078}T$ | V360L |
| | *mdoH* | 2544 | $T_{2055}C$ | W669R |
| | *araD* | 696 | $A_{23}G$ | Q8R |
| V25 | *wecC* | 1263 | $C_{783}CCCTGGC$ | G261GPG |
| | *pgm* | 1641 | $G_{520}T$ | E174X |
| | *araD* | 696 | $A_{23}G$ | Q8R |
| V26 | *wecC* | 1263 | $C_{783}CCCTGGC$ | G261GPG |
| | *pgm* | 1641 | $G_{520}T$ | E174X |
| | *araD* | 696 | $A_{23}G$ | Q8R |
| V27 | *wecC* | 1263 | $C_{783}CCCTGGC$ | G261GPG |
| | *pgm* | 1641 | $G_{520}T$ | E174X |
| | *araD* | 696 | $A_{23}G$ | Q8R |

*V25, V26, and V27 are isolated from the same plate and are likely derived from one clone

**Table S2 Bacterial strains used in this study.**

| Strains | Relevant genotype | References | Used in figures |
|---|---|---|---|
| MC4100 | [*F⁻ araD139 Δ(argF-lac) U169 rpsL150 relA1 flbB5301 ptsF25 deoC1 ptsF25 thi*] | Casadaban, 1976 | 1; 2; S2; S3; S5; S7 |
| BW25113 | *F- Δ(araD-araB)567 ΔlacZ4787::rrnB-3 λ- rph-1 Δ(rhaDrhaB)568 hsdR514* | Datsenko & Wanner, 2000 | S1 |
| NovaBlue | *endA1 hsdR17 (rK12– mK12+) supE44 thi-1 recA1 gyrA96 relA1 lac F' [proA+B+lacIqZΔM15::Tn10]* | Novagen | |
| NR754 | MC4100 *araD*$^{+}$ | Ruiz et al., 2008 | |
| RS101 | BW25113 Δ*tolQ::kan* | Shrivastava et al., 2017 | |
| RS102 | BW25113 Δ*tolA::kan* | Shrivastava et al., 2017 | |
| JW5100 | BW25113 Δ*tolB::kan* | Baba et al., 2006 | |
| TR51 | MC4100 *cpxR::spec* | Raivio et al., 1999 | |
| RS103 | BW25113 Δ*tol-pal::kan*# (During construction, the stop codon of the preceding orf *ybgC* was deleted by accident, resulting in the extension of YbgC by 10 aa) | This study | S1 |
| RS107 | BW25113 Δ*wecC::cm* | This study | |
| RS111 | BW25113 Δ*wecA::cm* | This study | |
| RS112 | BW25113 Δ*wecB::cm* | This study | |
| RS114 | BW25113 Δ*wecE::cm* | This study | |
| RS115 | BW25113 Δ*wecF::cm* | This study | |
| RS116 | BW25113 Δ*wecG::cm* | This study | |
| JXE370 | MG1655 Δ*wcaJ::kan* | Kevin Young | |
| TWB027 | BW25113 Δ*wecA::kan* | This study | |
| TWB028 | BW25113 Δ*wzxE::kan* | This study | |
| TWB029 | BW25113 Δ*wzzE::kan* | This study | |
| TWB030 | BW25113 Δ(*wecA-wzzE*)*::kan* | This study | |

| | | | |
|---|---|---|---|
| TWB031 | BW25113 Δ*wzyE::kan* pBAD33*wzyE* | This study | |
| TWB032 | BW25113 Δ*wzxB::kan* | This study | |
| RS174 | EH150 Δ*tolA::kan* | Shrivastava et al., 2017 | |
| RS280 | EH150 Δ*tolA::FRT* | Shrivastava et al., 2017 | |
| RS119 | MC4100 Δ*tolQ::kan* | Shrivastava et al., 2017 | 1; S2 |
| RS121 | MC4100 Δ*tolA::kan* | Shrivastava et al., 2017 | 1; 2; S2 |
| RS122 | MC4100 Δ*tolB::kan* | Shrivastava et al., 2017 | 1; S2 |
| RS273 | MC4100 *wecC** | This study | 1; S2 |
| RS139 | MC4100 Δ*wecC::cm* | This study | 1; 2; S3 |
| RS144 | MC4100 Δ*wecB::cm* | This study | 1 |
| RS147 | MC4100 Δ*wecF::cm* | This study | 1 |
| JXE158 | MC4100 Δ*tolQ::kan wecC** | This study | 1; S2 |
| JXE153 | MC4100 Δ*tolA::kan wecC** | This study | 1; S2 |
| JXE154 | MC4100 Δ*tolB::kan wecC** | This study | 1; S2 |
| JXE225 | MC4100 Δ*tolQ::kan* Δ*wecC::cm* | This study | 1 |
| JXE218 | MC4100 Δ*tolA::kan* Δ*wecC::cm* | This study | 1; 2; S5 |
| JXE334 | MC4100 Δ*tolB::kan* Δ*wecC::cm* | This study | 1B |
| JXE506 | MC4100 Δ*tolA::kan* Δ*wecB::cm* | This study | 1D |
| JXE244 | MC4100 Δ*tolA::kan* Δ*wecF::cm* | This study | 1D |
| JXE165 | MC4100 Δ*rcsC::cm* | This study | 2; S5 |
| JXE199 | MC4100 Δ*tolA::FRT* | This study | 2; S5 |
| JXE162 | MC4100 Δ*wecC::FRT* | This study | 2; S5 |
| JXE180 | MC4100 Δ*rcsC::cm* Δ*wecC::FRT* | This study | 2; S5 |
| JXE214 | MC4100 Δ*rcsC::cm* Δ*tolA::kan* | This study | 2; S5 |
| JXE562 | MC4100 Δ*rcsC::cm* Δ*tolA::FRT* Δ*wecC::FRT* | This study | 2; S5 |
| JXE375 | MC4100 Δ*wcaJ::kan* | This study | 2; S5; 6, 7 |
| JXE377 | MC4100 Δ*wcaJ::kan* Δ*tolA::FRT* | This study | 2; S5; 6, 7 |

| | | | |
|---|---|---|---|
| JXE539 | MC4100 Δ*wcaJ::kan* Δ*tolA::FRT* Δ*wecC::FRT* | This study | 2; S5 |
| JXE534 | MC4100 Δ*wcaJ::kan* Δ*wecC::cm* | This study | 2; S5; 7 |
| JXE538 | MC4100 Δ*wcaJ::kan* Δ*wecG::cm* | This study | 6, 7 |
| JXE523 | MC4100 Δ*wcaJ::kan* Δ*tolA::FRT* Δ*wecC::cm* | This study | 7 |
| JXE548 | MC4100 Δ*wcaJ::kan* Δ*tolA::FRT* Δ*wecG::cm* | This study | 6, 7 |
| TWB001 | MC4100 Δ*wcaJ::FRT* | This study | 2; 3; 4; 5; S4; S6; S8; S9; S10 |
| TWB002 | MC4100 Δ*wcaJ::FRT* Δ*tolA::FRT* | This study | 2; 3; 4; 5; S4; S6; S8; S9; S10 |
| TWB033 | MC4100 Δ*wcaJ::FRT* Δ*wecA::kan* | This study | 4; 5; S10 |
| TWB003 | MC4100 Δ*wcaJ::FRT* Δ*tolA::FRT* Δ*wecA::kan* | This study | 4; 5; S10 |
| TWB004 | MC4100 Δ*wcaJ::FRT* Δ*wecC::cm* | This study | 2; 3; 4; S4; S8; S9; S10 |
| TWB005 | MC4100 Δ*wcaJ::FRT* Δ*tolA::FRT* Δ*wecC::cm* | This study | 2; 3; 4; S4; S6; S8; S9; S10 |
| TWB006 | MC4100 Δ*wcaJ::FRT* Δ*wecC::cm* Δ*wecA::kan* | This study | 4 |
| TWB007 | MC4100 Δ*wcaJ::FRT* Δ*tolA::FRT* Δ*wecC::cm* Δ*wecA::kan* | This study | 4; S6 |
| TWB008 | MC4100 Δ*wcaJ::FRT* Δ*wecE::cm* | This study | 4; S10 |
| TWB009 | MC4100 Δ*wcaJ::FRT* Δ*tolA::FRT* Δ*wecE::cm* | This study | 4; S6; S10 |
| TWB010 | MC4100 Δ*wcaJ::FRT* Δ*wecE::cm* Δ*wecA::kan* | This study | 4 |
| TWB011 | MC4100 Δ*wcaJ::FRT* Δ*tolA::FRT* Δ*wecE::cm* Δ*wecA::kan* | This study | 4; S6 |
| TWB012 | MC4100 Δ*wcaJ::FRT* Δ*wecG::cm* | This study | 4 |
| TWB013 | MC4100 Δ*wcaJ::FRT* Δ*tolA::FRT* Δ*wecG::cm* | This study | 4; S6 |

| | | | |
|---|---|---|---|
| TWB014 | MC4100 Δ*wcaJ::FRT* Δ*wecG::cm* Δ*wecA::kan* | This study | 4 |
| TWB015 | MC4100 Δ*wcaJ::FRT* Δ*tolA::FRT* Δ*wecG::cm* Δ*wecA::kan* | This study | 4; S6 |
| TWB034 | NR754 Δ*wcaJ::FRT* Δ*wzxE::FRT* pBAD33*wzxE* | This study | S7 |
| TWB035 | NR754 Δ*wcaJ::FRT* Δ*wzxE::FRT* Δ*wzxB::kan* pBAD33*wzxE* | This study | S7 |
| TWB036 | NR754 Δ*wcaJ::FRT* pBAD33*wzyE* | This study | S7 |
| TWB037 | NR754 Δ*wcaJ::FRT* Δ*wzyE::kan* pBAD33*wzyE* | This study | 5, S7 |
| TWB025 | MC4100 Δ*wcaJ::FRT* Δ*wzzE::kan* | This study | 5, S10 |
| TWB026 | MC4100 Δ*wcaJ::FRT* Δ*tolA::FRT* Δ*wzzE::kan* | This study | 5, S10 |
| TWB038 | MC4100 Δ*wcaJ::FRT* Δ(*wecA-wzzE*)*::kan* | This study | 5 |
| TWB039 | MC4100 Δ*wcaJ::FRT* Δ*tolA::FRT* Δ(*wecA-wzzE*)*::kan* | This study | 5 |
| TWB040 | MC4100 Δ*wcaJ::FRT cpxR::spec* | This study | S10 |
| TWB041 | MC4100 Δ*wcaJ::FRT cpxR::spec* Δ*wzzE::kan* | This study | S10 |
| TWB042 | MC4100 Δ*wcaJ::FRT cpxR::spec* Δ*tolA::FRT* | This study | S10 |
| TWB043 | MC4100 Δ*wcaJ::FRT cpxR::spec* Δ*tolA::FRT* Δ*wzzE::kan* | This study | S10 |
| TWB044 | MC4100 Δ*wcaJ::FRT* Δ*waaL::kan* | This study | S8 |
| TWB045 | MC4100 Δ*wcaJ::FRT* Δ*tolA::FRT* Δ*waaL::kan* | This study | S8 |
| TWB046 | MC4100 Δ*wcaJ::FRT* Δ*wecC::cm* Δ*waaL::kan* | This study | S8 |
| TWB047 | MC4100 Δ*wcaJ::FRT* Δ*tolA::FRT* Δ*wecC::cm* Δ*waaL::kan* | This study | S8 |
| TWB048 | MC4100 Δ*wcaJ::FRT* Δ*wzxE::kan* | This study | S7 |
| TWB049 | MC4100 Δ*wcaJ::FRT* Δ*tolA::FRT* Δ*wzxE::kan* | This study | S7 |
| NR698 | MC4100 *lptD4213* | Ruiz et al., 2006 | S3 |
| TWB050 | MC4100 *lptD4213 ΔwecC::cm* | This study | S3 |
| TWB051 | MC4100 *bepA::*Tn5 | This study | S3 |
| TWB052 | MC4100 *bepA::*Tn5 *ΔwecC::cm* | This study | S3 |

**Table S3 Plasmids used in this study.**

| Plasmids | Relevant genotype/Description[a] | References |
|---|---|---|
| pKD3 | *oriR*$_{\gamma R6k}$ *bla FRT::cam::FRT* | Datsenko & Wanner, 2000 |
| pKD4 | *oriR*$_{\gamma R6k}$ *bla FRT::kan::FRT* | Datsenko & Wanner, 2000 |
| pKD13 | *oriR*$_{\gamma R6k}$ *bla FRT::kan::FRT* | Datsenko & Wanner, 2000 |
| pKD46 | *repA101*(ts) *oriR101 bla P*$_{araB}$ - (*gam bet exo*) | Datsenko & Wanner, 2000 |
| pCP20 | *λcI857*(ts) *repA101*(ts) *oriR101 bla cat λp*$_{R}$-FLP | Cherepanov & Wackernagel, 1995 |
| pKD4*tse2* | *oriR*$_{\gamma R6k}$ *bla FRT::kan*-P$_{rhaB}$-*tse2::FRT* | Khetrapal et al., 2015 |
| pTrc99a-*wecC* | Express *wecC* under IPTG inducible promoter; pBR ori; AmpR | This study |
| pTrc99a-*wecE* | Express *wecE* under IPTG inducible promoter; pBR ori; AmpR | This study |
| pTrc99a-*wecG* | Express *wecG* under IPTG inducible promoter; pBR ori; AmpR | This study |
| pTrc99a-*wecA* | Express *wecA* under IPTG inducible promoter; pBR ori; AmpR | This study |
| pDSW204*murA* | Express *murA* under IPTG inducible promoter; pBR ori; Amp$^R$ | Jorgenson et al., 2016 |
| pDSW204*uppS* | Express *uppS* under IPTG inducible promoter; pBR ori; Amp$^R$ | Jorgenson et al., 2016 |
| pBAD33*wzxE* | Express *wzxE* under control of arabinose inducible promoter P$_{ara}$; pACYC origin of replication; Cam$^R$ | This study |
| pBAD33*wzyE* | Express *wzyE* under control of arabinose inducible promoter P$_{ara}$ ; pACYC origin of replication; Cam$^R$ | This study |
| p*rpoH*P3::*gfp* | Express *gfp* under control of the *rpoH*P3 promoter; pBR322 origin of replication; Amp$^R$ | This study |
| p*gfp* | Same construct as p*rpoH*P3::*gfp* but without the *rpoH*P3 promoter (promoter-less); pBR322 origin of replication; Amp$^R$ | This study |

**Table S4 List of oligonucleotides used in this study.**

| Primer name | Sequence (5'-3')[b] |
|---|---|
| *wecC*-pKD4*tse2*-del-F | GCATTCTGGAAGCGTTAAAAAATAATCGGATATCACTATGATTGTGTAGGCTGGAGCTGC |
| *wecC*-pKD4*tse2*-del-R | GTTATCAGAATTTTTCTCATCAGCGCCAGACTCCTTTGGCCCATATGAATATCCTCCTTA |
| *wecA*-del-F | ATACTTCTGCTAATAATTTTCTCTGAGAGCATGCATTGTGGTGTAGGCTGGAGCTGCTTC |
| *wecA*-del-R | TGTGTCATCACATCCTCATTTATTTGGTTAAATTGGGGCTCATATGAATATCCTCCTTAG |
| *wecB*-del-F | CGCAAAGGCGCTCGCCGCTTATTCGAAGAGAATCGATGTGGTGTAGGCTGGAGCTGCTTC |
| *wecB*-del-R | ACAGAAATGGTCGCAAAACTCATAGTGATATCCGATTATTCATATGAATATCCTCCTTAG |
| *wecC*-del-F | GCATTCTGGAAGCGTTAAAAAATAATCGGATATCACTATGATTGTGTAGGCTGGAGCTGC |
| *wecC*-del-R | GTTATCAGAATTTTTCTCATCAGCGCCAGACTCCTTTGGCCCATATGAATATCCTCCTTA |
| *wecE*-del-F | GTAGAAAGCACCGCGTACTGGTTATACAGGTGATCACATGGTGTAGGCTGGAGCTGCTTC |
| *wecE*-del-R | CGCTTTTGCCAACGACATATCAGGAAAAGTAGTTCAACAACATATGAATATCCTCCTTAG |
| *wecF*-del-F | TTGTTGTGGCGTGTTTTTACTCTGGCGTAGGCGGGCATGAGTGTAGGCTGGAGCTGCTTC |
| *wecF*-del-R | CCACTGAATTGCAGCAGACTCATGCGACCTCCCTGGCGGCCATATGAATATCCTCCTTAG |
| *wecG*-del-F | CAAAATCATCGCTCCGGACGCAGGTTGAAGGATAACAATGGTGTAGGCTGGAGCTGCTTC |
| *wecG*-del-R | CTTTACAAAGAGAGGAAAATCATAGGTTGCCGGTGTAGTGCATATGAATATCCTCCTTAG |
| *rcsC*pkD3del-F | ACACTCCATCGGTCACCTGAGGCAGGAGCTTCGCCCCTTTGGTGTAGGCTGGAGCTGCTTC |
| *rcsC*pkD3del-R | CGTAAACGCCTTATCCGTCCTACGAATCCCGCGATTTCCTCATATGAATATCCTCCTTAG |
| *wzxE*KO5 | ACGGTAATTGCGACTTTGTTGAACTACTTTTCCTGATATGATTCCGGGGATCCGTCGACC |
| *wzxE*KO3 | AGTACGTGAATCAGTACAGTCATGCCCGCCTACGCCAGAGTGTAGGCTGGAGCTGCTTCG |
| *wecA*KO5 | ATACTTCTGCTAATAATTTTCTCTGAGAGCATGCATTGTGATTCCGGGGATCCGTCGACC |
| *wecA*KO3 | TGTGTCATCACATCCTCATTTATTTGGTTAAATTGGGGCTTGTAGGCTGGAGCTGCTTCG |

| | |
|---|---|
| *wzzE*KO5 | GTGGCAGCCCCAATTTAACCAAATAAATGAGGATGTGATGATTCCGGGGATCCGTCGACC |
| *wzzE*KO3 | GCTCACCGCAGCAGTGTTGCTATTTCGAGCAACGGCGGGTTGTAGGCTGGAGCTGCTTCG |
| *wzyE*KO5 | GGCAGCGGGCGTTGGCGATTGCCGCCAGGGAGGTCGCATGATTCCGGGGATCCGTCGACC |
| *wzyE*KO3 | GTGGTGTTGTTATTCATTGTTATCCTTCAACCTGCGTCCGTGTAGGCTGGAGCTGCTTCG |
| *wzxB*KO5 | TTACGTTAGATGAGCTTATCAGATTAAAATTAATTGCATGATTCCGGGGATCCGTCGACC |
| *wzxB*KO3 | ATAATCGTACATAAAATCCTCAGCAAACCAGTAATTTATTTGTAGGCTGGAGCTGCTTCG |
| *waaL*KO5 | AGTTTTGGAAAAGTTATCATCATTATAAAGGTAAAACATGATTCCGGGGATCCGTCGACC |
| *waaL*KO3 | TTCTTAAACTTGTTTATTCTTAATTAATTGTATTGTTACGTGTAGGCTGGAGCTGCTTCG |
| EcoRIwecA5 | ACAATGAATTCgtgAATTTACTGACAGTGAG |
| XbaIwecA3 | TACTGTCTAGAttaTTTGGTTAAATTGGGGC |
| wecC-N-NcoI | CAGTCCATGGGGAGTTTTGCGACCATT |
| wecC-C-XbaI | ACTCTAGATCAGCGCCAGACTCCTTTGG |
| wecE-N-NcoI | CATGCCATGGGGATTCCATTTAACGCACC |
| wecE-C-XbaI | GCTCTAGATCAGGAAAAGTAGTTCAACAAAG |
| wecG-N-NcoI | CATGCCATGGGGAATAACAACACCACGGCA |
| wecG-C-XboI | ACTCTAGATCATAGGTTGCCGGTGTAGTG |
| SacIRBSWzyE5 | ACAATGAGCTCAGGGAGGTCGCATGAGTCTG |
| XbaIWzyE3 | TACTGTCTAGATTATCCTTCAACCTGCGTCC |
| SacIRBSWzxE5 | ACAATGAGCTCAGGAGGTTTTCCTGATATGTCGTTGG |
| XbaIWzxE3 | TACTGTCTAGATCATGCCCGCCTACGCCAGA |

[b] restriction sites are underlined.
